# The human neuroblastoma SH-SY5Y cell line as a model to assess β-amyloid neurotoxicity: A systematic review and meta-analysis

**DOI:** 10.1101/2025.10.20.683497

**Authors:** Nathalia R. Pinheiro, Clarissa F. D. Carneiro, Giulia S. Cancelliero, Giovanna O. Nogueira, Glaucia M. Almeida, Nathalia Fernandes, Ana Paula Wasilewska-Sampaio, Samantha M. Martins, Antonio Felix, Olavo B. Amaral, Adriano Sebollela

**Author notes:** Corresponding authors:* Adriano Sebollela, Olavo B. Amaral. These authors contributed equally to this work (Nathalia R. Pinheiro, Clarissa F. D. Carneiro, Giulia S. Cancelliero). present affiliation: MUSC, USA.

## Abstract

The SH-SY5Y human neuroblastoma cell line is widely used as an *in vitro* model of β-amyloid (Aβ) neurotoxicity in Alzheimer’s disease (AD). However, the lack of standardized protocols for assessing Aβ toxicity - including differentiation strategies for SH-SY5Y cells - limits the comparability of results across studies. To address these issues, we conducted a systematic review and meta-analysis to evaluate how methodological factors influence Aβ-induced toxicity in SH-SY5Y cells. We included 359 eligible studies encompassing 1,192 MTT-based comparisons of cell viability between Aβ-treated and control SH-SY5Y cells. A three-level meta-analysis estimated mean cell viability after Aβ exposure at 63% of control levels (95% CI [61.6; 64.3]), with very high heterogeneity (I²=99.6%). Meta-regression identified significant associations between increased toxicity and higher Aβ concentrations, longer exposure durations, and the use of peptide preparations described as fibrils. Conversely, differentiation protocols, duration, and cell density did not significantly influence toxicity outcomes. Reporting quality was often poor, with frequent omissions regarding cell line origin, authentication, contamination testing, Aβ preparation details and nature of the experimental unit. Overall, our findings show robust Aβ toxicity in SH-SY5Y cells, primarily driven by dose, exposure time, and Aβ aggregation state, but not cell differentiation status. Our conclusions highlight the critical need for better reporting of Aβ exposure parameters to enhance reproducibility and translational potential in AD research.

## Introduction

Cognitive impairment in Alzheimer’s disease (AD) is widely attributed to synaptic dysfunction induced by aggregates of the β-amyloid peptide (Aβ) (Cline et al., 2018; Hardy & Selkoe, 2002; Sehar et al., 2022). Enzymatic cleavage of the amyloid precursor protein (APP) on the extracellular surface of neurons by BACE-1 and γ-secretase generates different isoforms of the Aβ peptide, with the most abundant isoforms having 40 (Aβ40) or 42 amino acid residues (Aβ42) (Querfurth & LaFerla, 2010). These peptides can assemble into structurally diverse aggregates with varying degrees of solubility and toxicity (Benilova et al., 2012; De et al., 2019; Michno et al., 2021; Sebollela et al., 2014; Velasco et al., 2012). Despite substantial efforts to characterize the neurotoxic properties of different Aβ species, progress has been hindered by inconsistencies across experimental models, Aβ concentrations, exposure durations, and aggregation protocols (Benilova et al., 2012; Fontana et al., 2020; Jan et al., 2010). This lack of standardization contributes to the limited generalizability of preclinical findings and may partly explain the high failure rates of Aβ-targeting therapies in clinical trials (Cummings et al., 2014; Mehta et al., 2017; Sehar et al., 2022).

The SH-SY5Y cell line is a widely used model in neuroscience research, including studies on AD (Agholme et al., 2010; De Medeiros et al., 2019; Fontana et al., 2020; Forster et al., 2016). Generated through multiple rounds of subcloning and selection from the parental SK-N-SH human neuroblastoma cell line, SH-SY5Y cells can be employed either in their undifferentiated state or following differentiation into a mature neuron-like phenotype, with all-trans retinoic acid (ATRA) being the preferential agent for this purpose (Biedler et al., 1978; Kovalevich & Langford, 2013). ATRA acts through the inhibition of cellular proliferation accompanied by increased expression of β-tubulin III, a cytoskeletal protein typical of mature neurons (Agholme et al., 2010; Encinas et al., 2000) and favours the emergence of a cholinergic phenotype (Kovalevich & Langford, 2013). When associated with brain-derived neurotrophic factor (BDNF), ATRA further enhances neuronal characteristics, including the formation of synaptic structures and axonal vesicle transport (Agholme et al., 2010; Krishtal et al., 2017). Alternative differentiation protocols have also been explored, including the use of specialized media (e.g., N2) or the addition of alternative growth factors (e.g., FGF) (Pedersen et al., 2021).

Several studies, including those using the SH-SY5Y cell line (Krishtal et al., 2017), have proposed that toxicity mechanisms elicited by Aβ aggregates involve direct interactions with synaptic structures (Alfonso et al., 2014; Lacor et al., 2007). This raises concerns about the suitability of undifferentiated SH-SY5Y cells, which do not present extended neurites or functional synapses, as a model for studying Aβ-induced toxicity. Moreover, SH-SY5Y cells differentiated using distinct culture media exhibit differential toxicity responses to Aβ42 aggregates exposure (Krishtal et al., 2019), suggesting that the differentiation mechanistic route and final phenotype distribution of the cell culture impact toxicity outcomes. Such differences may explain the wide range of Aβ concentrations used in studies using SH-SY5Y as a model - including some that are much higher than those found *in vivo* (Hu et al., 2009; Seubert et al., 1992). The combination of variability in differentiation protocols, Aβ aggregation preparation, and Aβ concentrations is likely to affect cellular responses and contribute to poor commensurability across studies (De Medeiros et al., 2019; Krishtal et al., 2019).

Motivated by the widespread use of SH-SY5Y cells as a model to investigate Aβ toxicity, we conducted a systematic review and meta-analysis to determine the main factors influencing Aβ toxicity in this cell line. Specifically, we examined parameters related to Aβ exposure - including aggregation state, concentration, and duration - as well as cell line variables such as differentiation protocol and cell density. In addition, we carried out a comprehensive assessment of methodological aspects and reporting quality in the included articles. Our results point out windows of opportunity for improving reporting standards and may offer practical guidance for optimizing the use of SH-SY5Y cells in future studies of Aβ neurotoxicity and AD pathology.

## Methods

### Search Strategy

The study protocol was preregistered at the Open Science Framework (https://doi.org/10.17605/OSF.IO/EV4TZ), and deviations from the preregistered protocol are described in https://osf.io/jc5bz. Initially, we created a list of 14 sentinel articles to guide the development of search strategies (Supplementary Table 1). Our final search string comprised two components, one with keywords related to the *in vitro* model and the other to AD or the β-amyloid peptide (Supplementary Table 2). We conducted this search among titles and abstracts of articles indexed in PubMed and EMBASE on December 18, 2020, filtering for those written in English. The total number of articles retrieved from each database is reported in Supplementary Table 2.

An additional search, using the same search strategy and search string, was conducted to analyze the number of studies published after the review ended. The abstracts of these studies were screened by querying a large-language model (GPT 5.2) in a structured fashion (https://osf.io/fqe85/overview), and a 10% sample was analyzed to assess the commensurability of article features with the initial sample.

### Title and abstract screening

Reviewers analyzed articles and removed duplicates using Rayyan (Ouzzani et al., 2016). Inclusion criteria were (i) articles written in English that (ii) presented original results, (iii) described experiments with administration of the β-amyloid peptide of synthetic origin or purified from a recombinant source, (iv) described experiments with SH-SY5Y cells as a model, and (v) were not retracted. The title and abstract screening step was carried out following the same criteria in the screening pilots, with each article evaluated by a single reviewer and 10% of articles undergoing double screening to calculate agreement levels. The agreement between reviewer pairs was measured by absolute percentage and Cohen’s kappa.

### Full-text screening

During full-text screening, each article included was assessed by a reviewer distinct from the one who had performed the initial title and abstract screening. Inclusion criteria were the same as in the previous step, with the addition of the following: (i) use of SH-SY5Y cells exposed to the Aβ peptide in its isoform(s) 1-37 to 1-43; (ii) use of a control group of SH-SY5Y cells, subject to the same culture conditions but not exposed to Aβ; (iii) use of toxicity outcomes measuring cell viability, cell death or oxidative stress markers; (iv) presentation of sample size, mean and standard deviation (SD) or standard error of the mean (SEM) for each group. If any of the required statistics were unclear or missing, the corresponding authors were contacted to request the necessary information. At this stage, the figure containing experiments that met the inclusion criteria was recorded using Airtable (https://airtable.com/).

### Data Extraction

Figure data were extracted by a reviewer distinct from those who included the article in the screening stages. Variables were collected at the article, figure/table, results, and protocol levels and encompassed article metadata, experimental design, results, and protocol variables such as control and treatment group outcomes and details on Aβ, cell culture and treatment conditions. The complete set of extracted variables is described in Supplementary Table 3. Gsys (GSYS 2.4.7) was used to extract values from the graphs and Airtable (https://www.airtable.com/) was used to record data.

### Data Processing

Data sheets with data extracted by different reviewers were combined using R (4.4.3) (Team, 2024). Extracted data for the description of experimental units was reviewed and categorized. Information on the number of plated cells per well and plate size was used to calculate cell density as a new variable. The description of medium supplements was reviewed, which led to the creation of two dichotomous variables to indicate the use of antibiotics and glutamine. After this manual cleaning step, data was imported back into R to conduct all subsequent cleaning, analysis, and visualizations. Comparisons with no data on either group’s means or sample size, as well as comparisons with missing variation for the treated group, were excluded from the analyses. When variability was reported as SEM, the SD was calculated as SEM×√sample size. If the meaning of error bars was unclear, they were assumed to represent SEM. If error bars were presented for the treated group only, we assumed that paired normalization was performed (with each treated unit divided by a particular control value).

### Data Synthesis

All analysis scripts and datasets are available at OSF (https://doi.org/10.17605/OSF.IO/EV4TZ). Packages used are listed in Supplementary Table 4 (Calcagno, 2020; Kossmeier et al., 2020; Viechtbauer, 2010; Wickham et al., 2019; Wickham & Bryan, 2023). All analyses were conducted with both log-transformed ratio of means (ROM/ROMC) and standardized mean difference (SMD) as effect size measures. When the control group had no SD or SEM reported, we considered it as zero and used measure = “ROMC” or “SMC” in the escalc function in metafor. Measure = “ROM” or “SMD” was used when SD or SEM was reported for the control group. For all ratio of means analyses, we transformed the resulting effect estimates to a linear scale (i.e., % of control) when displaying results by taking the natural exponential of the meta-analytical results.

Random-effects meta-analyses (two-level models) were calculated using the restricted maximum likelihood estimator for τ^2^. Based on these results, publication bias was assessed by trim-and-fill analyses using both R0 and L0 estimators, and by Egger’s regression using the inverse of SEM as the predictor variable. Three-level meta-analysis models were calculated using random effects at both the article and experiment levels.

Three-level univariate meta-regressions were performed using differentiation duration (no differentiation = 0 days), Aβ aggregation state, Aβ concentration, Aβ exposure duration and cell density. For meta-regressions using Aβ concentration as a moderator, we excluded comparisons in which Aβ concentrations were higher than 100 µM (18 comparisons), as high concentrations of this peptide in solution are prone to fast aggregation into insoluble species (Burdick et al., 1992). Three-level multivariate meta-regressions used the maximum likelihood estimator of τ^2^, and all combinations of variables from the selected list were tested in multivariable models. The best models were ranked by corrected Akaike Information Criteria (AICc). Additionally, we calculated the R^2^ for the model and performed a Q test of moderators for each variable (including all dummy variables for each categorical moderator) to obtain p-values for individual variables. We also built a multivariable model using Aβ concentration and Aβ exposure duration, including an interaction term to evaluate possible interactions between these two variables.

## Results

### Selection of experiments

The study selection process is illustrated in the PRISMA flowchart (Figure 1). A total of 9,391 articles were screened, resulting in 1,364 articles included for full-text screening. Mean agreement between reviewers in the sample of articles undergoing double screening was 86.6%, with a Cohen’s kappa of 0.67 (see https://osf.io/bg4qt). From the 1,364 articles analyzed in the full-text screening step, 553 were included for data extraction.

**Figure 1.**
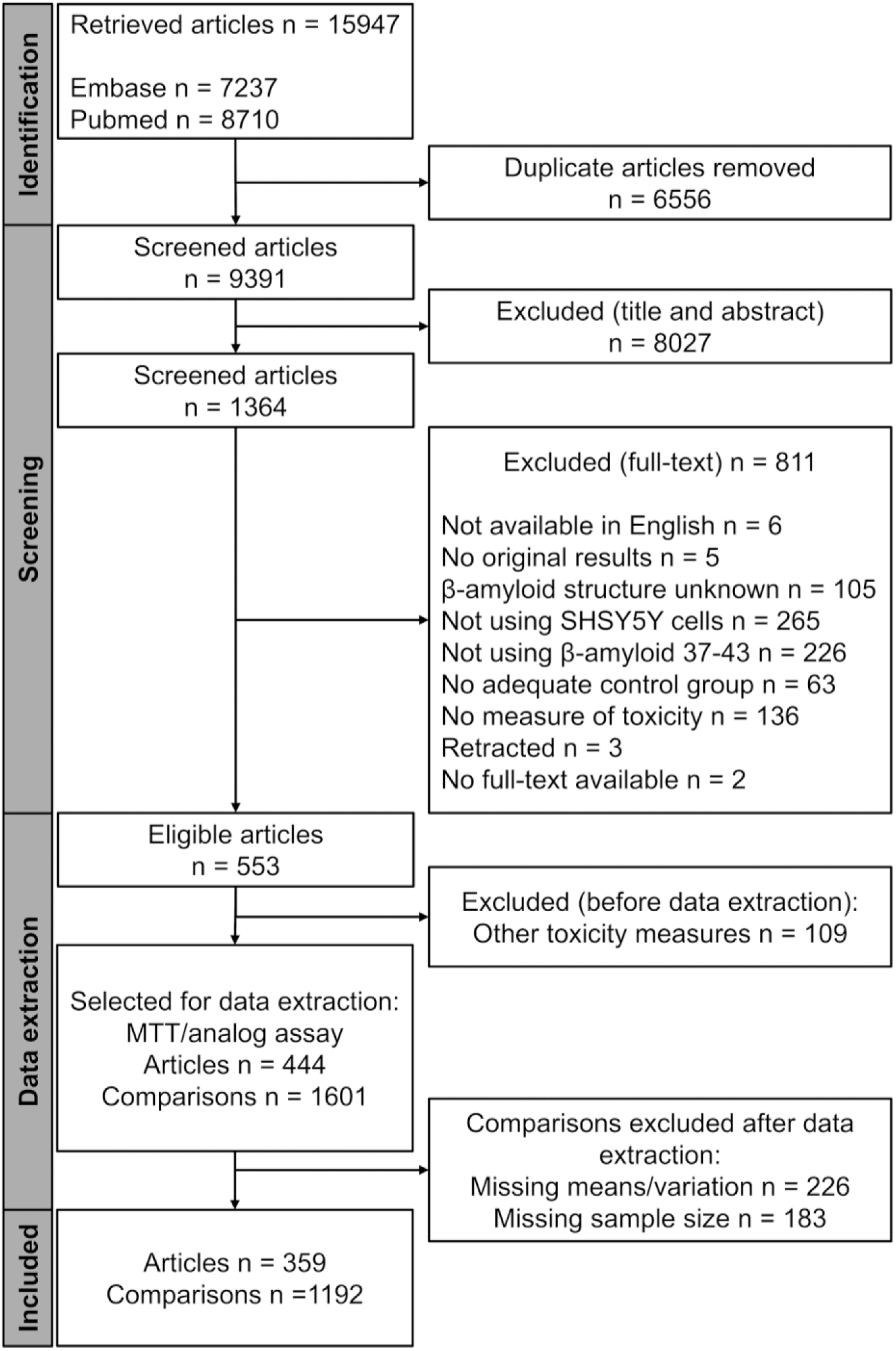
PRISMA flowchart of the systematic review. The flowchart summarizes the screening rounds and data extraction, which resulted in 359 articles, including 1,192 comparisons between control and treatment groups, selected for data analysis.

After the second screening, an initial analysis of the toxicity readout methods (Supplementary Table 5) revealed metabolic-based viability assays to be the most common readout, with 1,386 comparisons (54% of total). MTT was the most widely used reagent in these assays (>80% of comparisons; Supplementary Table 6). Based on the widespread use of these assays, we opted to extract data only from these comparisons for feasibility reasons. During data extraction, 215 additional comparisons in the selected articles were identified as eligible by reviewers, which led to a total of 444 articles and 1,601 comparisons. Of these, 226 comparisons were excluded due to missing information on means or variability, and 183 were excluded due to lack of information on sample size, leaving 1,192 comparisons from 359 articles for analysis.

### Sample description

Included articles were published between 1998 and 2021 (Supplementary Figure 1) and presented between 1 and 52 comparisons each, with a median of 2 and a mean of 3.3 comparisons per article (Supplementary Figure 2). In most of the selected studies (69%), the assay was used to test whether a given intervention could reverse the toxicity elicited by Aβ. Cell origin was reported in only 55% of articles. Reported sources were mostly cell banks (51% of articles), predominantly from the American Type Culture Collection (ATCC) (30%) or the European Collection of Authenticated Cell Cultures (ECACC) (8%) (Supplementary Table 7). The most commonly used culture medium was a combination of DMEM and F12 (36%), followed by DMEM (31%). In 8.5% of comparisons, culture medium was not reported. Most experiments included antibiotics in the culture medium (72%), and approximately 40% of the experiments included glutamine supplementation. In most experiments, the cell culture medium was supplemented with 10% fetal bovine/calf serum (FBS/FCS), but a wide range of serum types and concentrations was observed (Supplementary Table 8).

Table 1 summarizes variables related to the Aβ exposure protocol. All experimental comparisons but one described a single exposure to Aβ. Aβ1-42 was the most used isoform (81% of comparisons), followed by Aβ1-40 (18%), which is expected given the implication of these two isoforms in AD (Glenner & Wong, 1984); (Walsh et al., 1997); (De et al., 2019). In most experiments, neither the source of the peptide nor the species’ identity was reported (52% and 80%, respectively). The oligomeric (soluble) form was the most prevalent aggregation status reported (38%), followed by fibrils (9%). Almost half of the experiments did not provide a clear description of the state of aggregation (45%) and likely used mixtures of different aggregates (soluble and insoluble). A clear description of the control (unexposed) condition (e.g., medium only, vehicle, etc.) used was missing in 44% of comparisons. Exposure duration and concentration of Aβ were reported in 99% of cases. A wide range of values was observed, with a mean (± S.D.) exposure duration of 33.2 h (± 16.9) and a mean Aβ concentration of 41.5 μM (± 875.4).

**Table 1.** Aβ exposure protocols. The percentage of each Aβ treatment protocol variable is related to the 1192 included comparisons. The number (N), mean, standard deviation (SD), median, minimum (min), and maximum (max) values of exposure duration and concentration variables are displayed.

| Treatment variable | Category | Comparisons |  |
| --- | --- | --- | --- |
|  |  | N | % |
| Aβ isoform | 1-42 | 963 | 80.8 |
|  | 1-40 | 217 | 18.2 |
|  | 1-43 | 10 | 0.8 |
|  | 1-38 | 2 | 0.2 |
| Aβ origin | Synthetic | 524 | 44.0 |
|  | Recombinant | 52 | 4.4 |
|  | Not reported | 616 | 51.7 |
| Aβ species identity | Human | 232 | 19.5 |
|  | Rat | 5 | 0.4 |
|  | Not reported | 955 | 80.1 |
| Aβ aggregation | Oligomers | 462 | 38.8 |
|  | Fibrils | 103 | 8.6 |
|  | Monomers | 92 | 7.7 |
|  | Not reported | 535 | 44.9 |
| Control description | Vehicle | 344 | 28.9 |
|  | Medium only | 286 | 24.0 |
|  | Other | 41 | 3.4 |
|  | Not reported | 521 | 43.7 |
| Treatment variable | Estimate | Data |  |
| Exposure duration (hours) | Reported | 99.1% |  |
|  | Mean (SD) | 33.2 (16.8) |  |
|  | Median (min, max) | 24 (0, 144) |  |
| Concentration (μM) | Reported | 98.6% | |
|  | Mean (SD) | 41.5 (875.4) |  |
| | Median (min, max) | 10 ($10^{-5}$, $3 \times 10^4$) | |

Variables related to the differentiation protocol are summarized in Table 2. SHSY-5Y cells underwent differentiation in 270 comparisons (23%). Culture medium composition used for differentiation was detailed less often than media used for standard cell culturing. As for routine culturing, DMEM and DMEM+F12 were the most frequently reported medium options. The medium was supplemented with FBS/FCS in 47% of comparisons, with 5% or less being the most common concentration range (32%). Regarding the differentiation method, ATRA was employed in 51.5% of comparisons, while ATRA plus (ATRA associated with another agent, such as BDNF, N2, or FGF) appeared in 30%. In almost 20% of comparisons with differentiated cells, the duration of differentiation was not reported, and a similar fraction did not describe the concentration of ATRA. Among comparisons describing these variables, mean differentiation duration was 6.6 (± 3.2) days, and mean ATRA concentration was 9.2 µM (± 3.4).

**Table 2.** Differentiation protocol variables. Numbers and percentages of comparisons for each category are presented. Percentages are relative to the 270 comparisons in which cells underwent differentiation. The number (N), mean, standard deviation (SD), median, and range of differentiation duration (days), and concentration of ATRA (µM) are also shown.

| Protocol variable | Category | Comparisons |  |
| --- | --- | --- | --- |
|  |  | N | % |
| Method | ATRA | 139 | 51.5 |
|  | ATRA plus | 80 | 29.6 |
|  | Other | 25 | 9.3 |
|  | Not reported | 26 | 9.6 |
| Serum type | FBS/FCS | 127 | 47 |
|  | No serum | 15 | 5.6 |
|  | Not reported | 128 | 47.4 |
| Serum concentration | 5% or less | 86 | 31.9 |
|  | 10% | 41 | 15.2 |
|  | No serum | 15 | 5.6 |
|  | Not reported | 128 | 47.4 |
| Medium | DMEM_F12 | 111 | 41.1 |
|  | DMEM | 38 | 14.1 |
|  | Other | 25 | 9.3 |
|  | Not reported | 96 | 35.6 |
| Antibiotics | Yes | 66 | 24.4 |
|  | Not reported | 204 | 75.6 |
| Glutamine | Yes | 38 | 14.1 |
|  | No | 21 | 7.8 |
|  | Not reported | 211 | 78.1 |
| Protocol variable | Estimate | Data |  |
| Differentiation duration (days) | Reported | 82.2% |  |
|  | Mean (SD) | 6.6 (3.2) |  |
|  | Median (min, max) | 6 (1, 14) |  |
| ATRA concentration (μM) | Reported | 80.7% | |
|  | Mean (SD) | 9.2 (3.4) |  |
|  | Median (min, max) | 10 (0, 20) |  |

### Reporting quality

Article-level variables related to reporting are presented in Table 3. None of the selected studies reported a sample size calculation, and protocol registration was reported in only 10 studies (2.8%). A conflict of interest statement was present in 51% of studies and was more frequent in recent articles (Supplementary Figure 3). In a significant proportion of articles (44.6%), the origin of the cells was not reported. Cell line authentication and testing for mycoplasma contamination were reported in only 2 and 3 articles, respectively, and protocols for these tests were not provided in any of them. Moreover, the meaning of sample sizes was not always clear. Although a generic description such as “independent experiments” or “replicates” was usually provided (ca. 64% of comparisons combined; Table 4), there is no guarantee that authors agree on the meaning of these terms. For instance, one may refer to “independent experiments” as either repeating the assay using a single culture batch, independent culture batches, or different passages of the same batch.

**Table 3.**
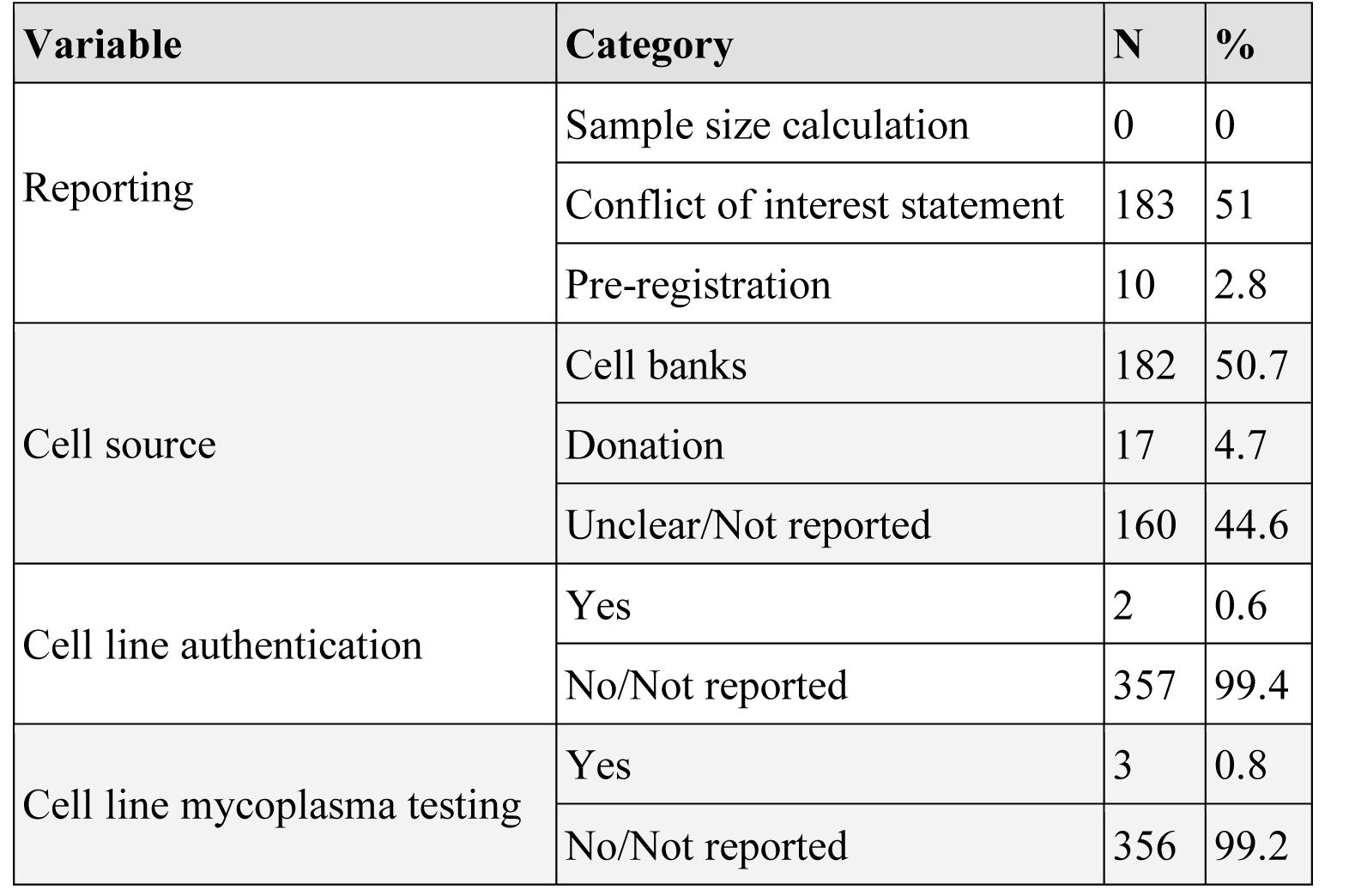
Article-level reporting and cell line quality control. Numbers (N) of articles and percentages (in relation to the 359 included articles) for each category are presented.

| Variable | Category | N | % |
| --- | --- | --- | --- |
| Reporting | Sample size calculation | 0 | 0 |
|  | Conflict of interest statement | 183 | 51 |
|  | Pre-registration | 10 | 2.8 |
| Cell source | Cell banks | 182 | 50.7 |
|  | Donation | 17 | 4.7 |
|  | Unclear/Not reported | 160 | 44.6 |
| Cell line authentication | Yes | 2 | 0.6 |
|  | No/Not reported | 357 | 99.4 |
| Cell line mycoplasma testing | Yes | 3 | 0.8 |
|  | No/Not reported | 356 | 99.2 |

**Table 4.** Experimental unit descriptions. “Others” included cell cultures, independent repetitions, independent replicates, independent runs, independent sets of studies, and observations.

| **N definition** | **N** | **%** |
| --- | --- | --- |
| Independent experiments | 542 | 45.5 |
| Replicates | 217 | 18.2 |
| Experiments | 108 | 9.1 |
| Wells | 24 | 2 |
| Assays | 19 | 1.6 |
| Independent determinations | 13 | 1.1 |
| Samples | 12 | 1 |
| Determinations | 8 | 0.7 |
| Independent experimental measurements | 6 | 0.5 |
| Biological replicates | 5 | 0.4 |
| Others | 9 | 0.8 |
| Not informed | 229 | 19.2 |

We also conducted a brief scoping review of the literature published between December 2020 and April 2025, applying the same search strategy, including the search string and databases (Supplementary Figure 4). After duplicate removal, 3,321 articles were retrieved. This set was subjected to an abstract screening using the same eligibility criteria with the support of an IA-based tool developed by our collaborators at BRISA (https://osf.io/fqe85/overview), resulting in the exclusion of 2,780 articles and the inclusion of 541. Given the substantial number of eligible articles and time constraints, we opted to perform full-text screening and qualitative data extraction on a random sample of 10% of these studies, resulting in the inclusion of 29 articles. Inclusion rates in both the abstract and full-text screening stages in these samples were similar to those obtained in the original set of articles.

Among the included articles, 76% (22) employed cell viability assays to assess Aβ-induced toxicity, a proportion similar to that observed in the original review (80%). Qualitative data from these 22 studies were collected (see Supplementary Table 9). Reporting of Aβ variables remained poor, with most studies lacking information on peptide origin, species, or aggregation status. Consistent with our previous findings, Aβ_42_ was the most frequently used sequence (82%), and oligomers were the most commonly reported aggregation state (32%). Neuronal differentiation was still uncommon, with only 3 out of 22 studies (14%) reporting the use of differentiated SH-SY5Y cells, compared to 23% in the original review.

### Main meta-analyses

We initially ran a random-effect meta-analysis including all 1,192 comparisons based on standardized mean differences as the effect size measure, as planned in our preregistered protocol. This model yielded an effect size (Hedges’ g) of -5.47 (95% CI [-5.7, -5.2], p <0.0001) (Figure 2A) and high heterogeneity (I^2^ = 93.77%, Q-test p value < 0.0001), indicating that the retrieved differences in effect size would be very unlikely to occur by sampling error alone.

**Figure 2.**
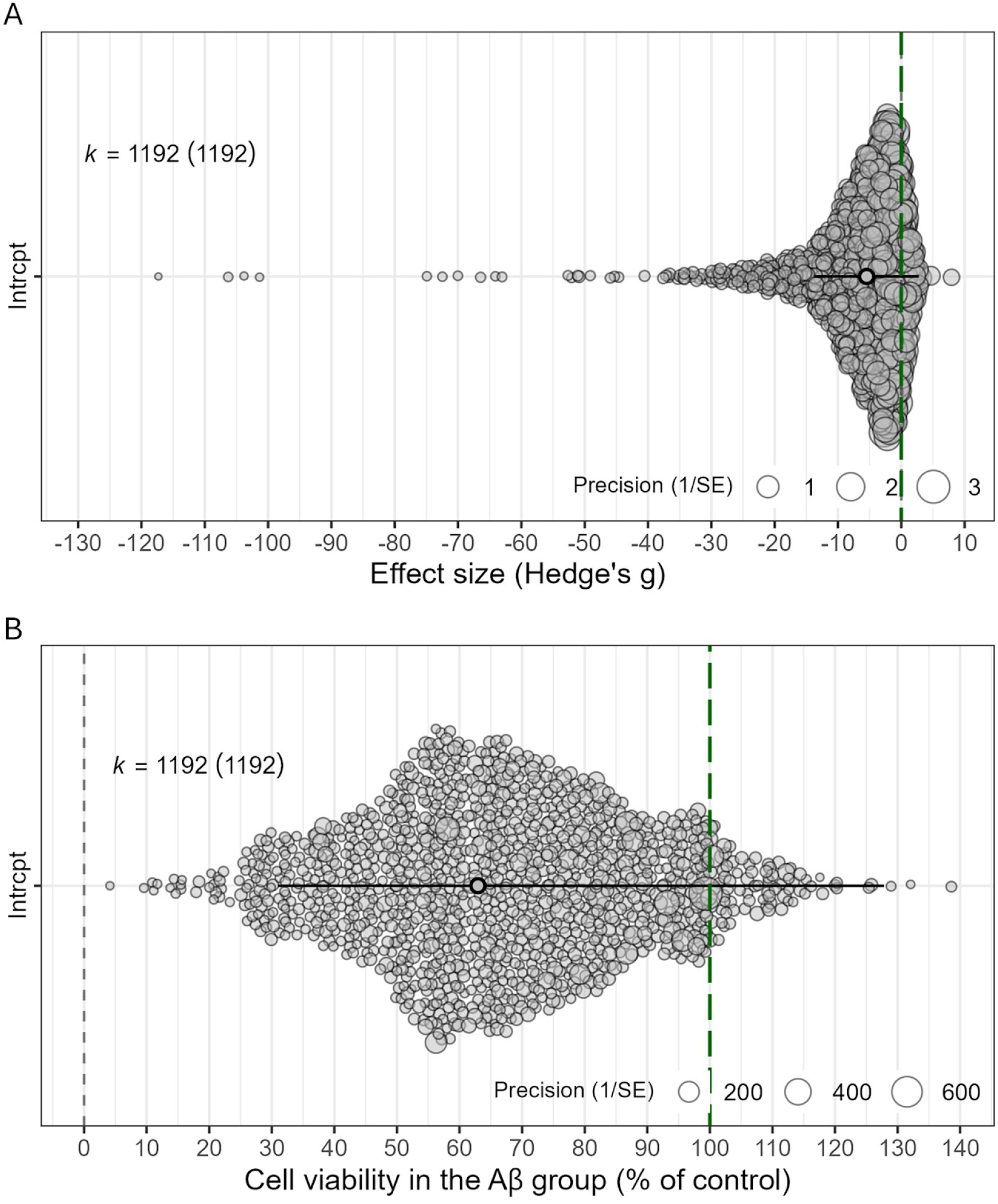
Orchard plots of two-level random-effects meta-analyses. Standardized effect sizes (A) or relative effect sizes expressed as ratios of means (B) results of two-level random-effects meta-analyses using all 1,192 comparisons included in this study. Each comparison is represented by a grey circle, with its position on the x-axis indicating the effect size (i.e., standardized difference between groups in A, % viability in the treated group relative to control in B) and its size indicating the inverse of standard errors. The green dashed line indicates the absence of effect (i.e., zero difference or 100% viability).

The use of standardized effect sizes, however, presented two problems. First, in a significant number of studies (51.6%), no error bars for the control group were presented (likely because each treated unit had its value normalized by a particular control unit). In these cases, standardized effect sizes cannot be accurately estimated, as pooling the error bars for both groups will not provide the true pooled standard deviation. Second, the uncertainties over the meaning of experimental units (Table 4) raised questions about whether the degree of independence between experimental units was commensurable between studies. This concern was further strengthened by the presence of studies reporting extremely small standard deviations between experimental units, leading to very large standardized effect sizes - potentially due to units not being truly independent from each other in each group (as in the case of technical replicates). Beyond that concern, the mere fact that error bars were very small led to unreliable estimates of SD or SEM from graphs (and thus to unreliable standardized effect sizes).

Because of this, we chose to use the ratio of means as our main effect size measure, as it (a) provides results that are commensurable between normalized and non-normalized data and (b) is not distorted by underestimation of variability (although studies with very small standard errors will still have greater weight in meta-analyses). Using this approach, the random-effects meta-analysis yielded a log ratio of means ([95% CI]) of -0.46 [-0.48,-0.44], p<0.0001, which corresponds to mean cell viability in the Aβ-treated group being 63% [61.6%, 64.3%] of that observed in controls (Figure 2B). Heterogeneity was still very high (I²=99.6%; Q-test p-value <0.0001). All our subsequent analyses are based on this measure, but alternative analyses using standardized effect sizes are presented as supplementary material.

### Publication Bias Analysis

Publication bias was assessed by funnel plot asymmetry (Figure 3) and Egger’s regression test (p=0.1587). Trim-and-fill analysis obtained with the R0 method showed strong evidence of publication bias (Supplementary Table 10), with 86 missing studies and an adjusted meta-analysis effect size of -0.40 [-0.42, -0.37], while the L0 method found no missing studies and thus yielded no difference from the main meta-analysis. The results observed in Figure 3 show that experiments in which viability in the treated group is above 100% are rarely reported. This is arguably expected, as in most experiments (∼70%), Aβ exposure was used with the purpose of testing for reversal of its toxic effect with another intervention. In these cases, it is likely that the absence of toxicity is interpreted as a methodological failure that precludes evaluation of the treatment and thus ends up unreported.

**Figure 3.**
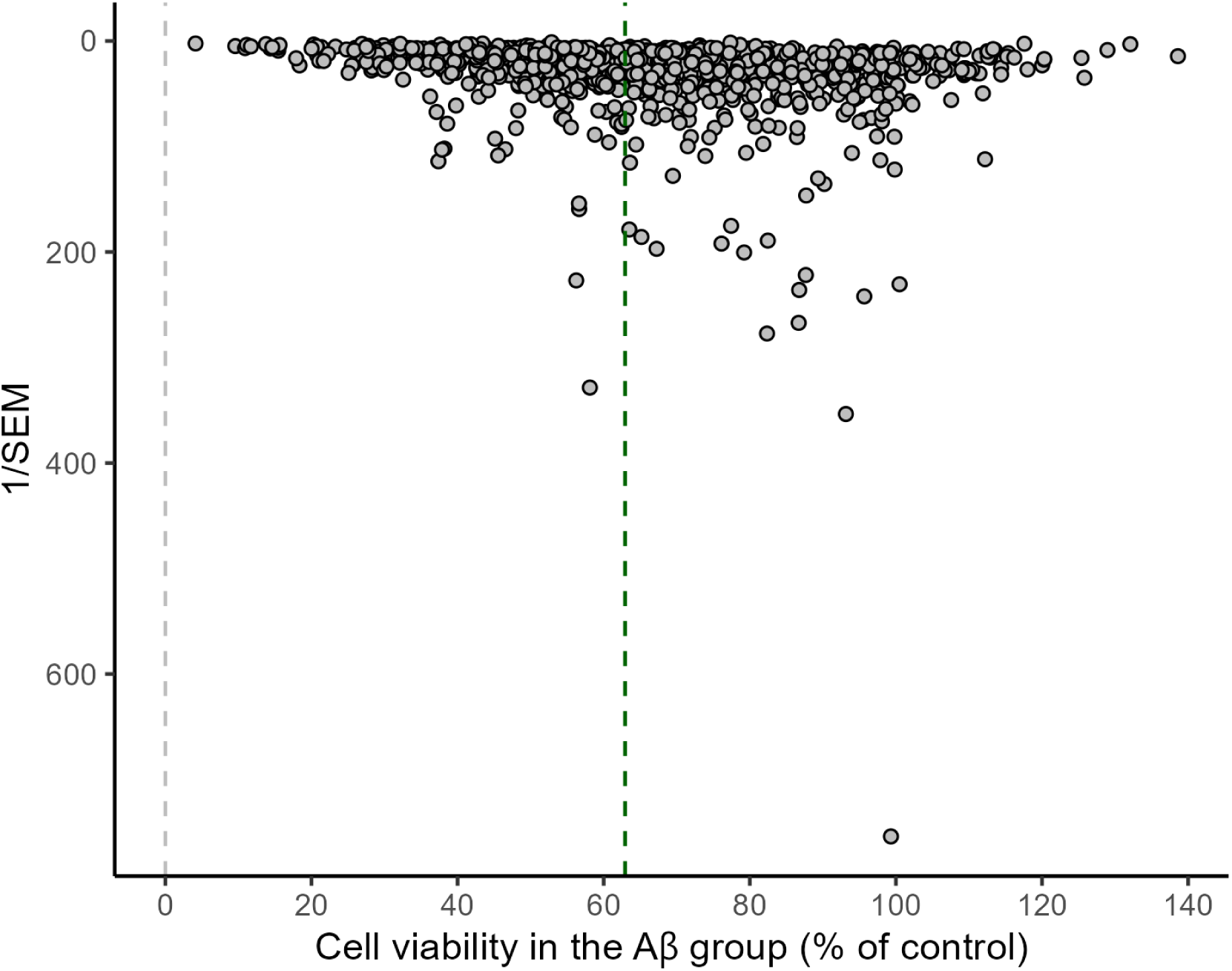
Funnel plot for all 1,192 comparisons included in this study. Each comparison is represented by a grey circle, with its position on the x-axis indicating the cell viability in the Aβ-treated group (% of control) and on the y-axis indicating the study precision (1/SEM). The overall effect is indicated by the dashed green line.

### Three-level meta-analyses

Given the nested structure of the data, with a mean of 3.3 comparisons per article, we conducted a three-level meta-analysis, using the experiment and study of origin as independent sources of variability nested within each other. We found the meta-analytical estimate to be similar to that of the two-level analysis, with a log ratio of means ([95% CI]) of -0.50 [-0.53, -0.47], p<0.0001, which corresponds to a mean cell viability in the Aβ-treated group of 61% [59%, 63%], p<0.0001. The variance ([95% CI]) was 0.09 [0.08, 0.1] with I^2^=66.3% between comparisons and 0.04 [0.03, 0.06] with I^2^=33.4% between articles, indicating that the heterogeneity between experiments within an article was larger than that between articles. This may reflect the use of different experimental conditions (e.g., Aβ concentrations, exposure duration, etc.) within individual articles, including some that may be expected to have no effect and are used as negative controls.

### Meta-regressions

The primary goal of our meta-analysis was not to obtain an effect estimate for the overall sample of articles, but rather to explore how different variables impact this effect. Before data collection, we selected the following variables to analyze in meta-regressions: differentiation method, differentiation duration, Aβ state of aggregation, Aβ concentration, Aβ exposure duration, cell density, and outcome measure; the latter was excluded as we only extracted data from cell viability assays. We also removed the differentiation method due to the limited number of comparisons in which a differentiation protocol was carried out. Analyses are based on all results for which the variable was reported and are shown in Table 5 and Figure 4 (with meta-regressions based on SMD available as Supplementary Table 11 and Supplementary Figure 5). Toxicity of Aβ was associated with higher concentrations, longer exposure, and Aβ preparations reported as fibrils or with no aggregation status reported (as compared to Aβ monomers). Differentiation duration showed a trend of association with lower toxicity, which was more significant when analysis was carried out using SMDs. Cell density, on the other hand, had no significant impact on effect sizes.

**Figure 4.**
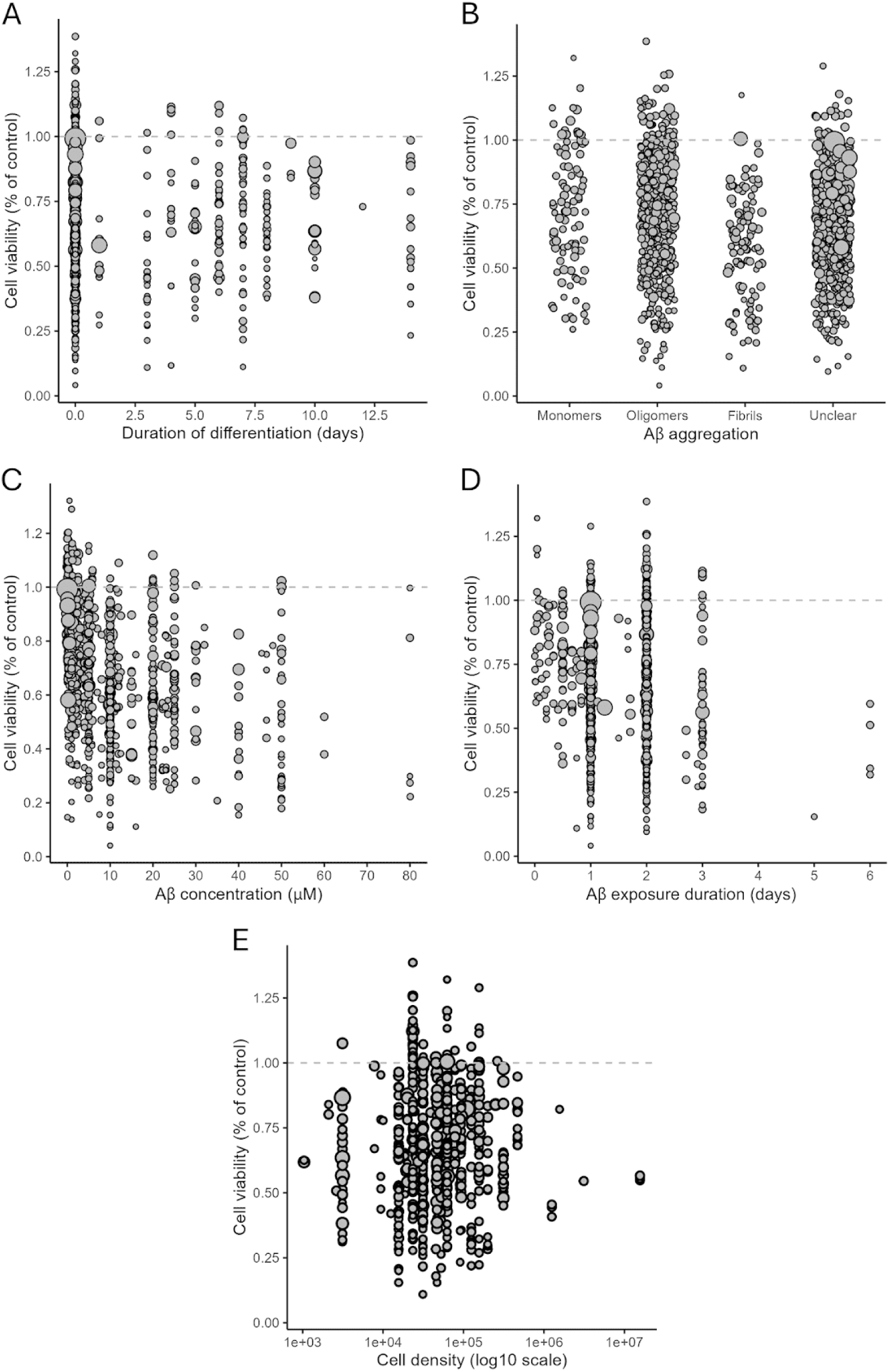
Univariate meta-regressions. Each experiment is represented by an open circle, with size corresponding to the inverse of variance (i.e., larger circles are more precise). Aβ concentrations above 100 µm were excluded from the concentration meta-regression.

**Table 5.** Three-level meta-regression models for protocol variables. Models have separate nested levels for articles and individual comparisons. Grey lines contain sample sizes, intercept effect sizes as ratio of means and cell viability, R^2^ for the moderator and Q-test p-values for the moderator. White lines contain betas indicating the additional contribution of each unit/category to the effect size (as well as sample sizes and p-values for individual categories). For categorical variables, reference groups are described in the first column, with the sample size for these groups indicated in parentheses in the second column. I^2^ and intercept p-values for all moderators were ∼99.6% and <0.0001, respectively. Moderator R^2^ values are calculated as the difference between total variances in the model with no moderators and in the tested model, divided by the total variance in the model with no moderators. Aβ concentrations above 100 µm were excluded from the concentration meta-regression.

| Moderators and categories | Sample size | Effect size [95% C.I.] | Cell Viability | R<sup>2</sup> | p-value |
| --- | --- | --- | --- | --- | --- |
| Duration of differentiation | 1144 | -0.51 [-0.54, -0.48] | 60% [58.3%, 61.9%] | 2.7% | 0.0595 |
| (day) |  | +0.01 [-0.0004, +0.02] | +1% [0%, +2%] |  |  |
| State of aggregation (reference=monomers) | 1192 (92) | -0.51 [-0.55, -0.47] | 60% [57.7%, 62.5%] | 2.2% | 0.0004 |
| Fibrils | 103 | -0.11 [-0.2, -0.02] | -10.4% [-18.1%, -2%] |  | 0.0215 |
| Oligomers | 462 | +0.05 [-0.01, +0.11] | +5.1% [-1, +11.6%] |  | 0.123 |
| Unclear | 535 | -0.13 [-0.24, -0.02] | -12.2% [-21.3%, -2%] |  | 0.016 |
| Aβ concentration | 1123 | -0.35 [-0.39, -0.32] | 70.5% [67.7%, 72.6%] | 15.7% | <0.0001 |
| (μM) | | -0.013 [-0.015, -0.011] | -1.3% [-1.5%, -1.1%] | | |
| Aβ exposure duration | 1180 | -0.35 [-0.41, -0.30] | 70.5% [66.4%, 74.1%] | 2.7% | < 0.0001 |
| (day) |  | -0.11 [-0.14, -0.07] | -10.4% [-13.1%, -6.8%] |  |  |
| Cell density | 690 | -0.50 [-0.54, -0.46] | 60.7% [58.3%, 63.1%] | 0% | 0.798 |
| 10<sup>3</sup> cells/cm<sup>2</sup> |  | -4.2x10<sup>-12</sup> [-3.6x10<sup>-12</sup>, 2.8x10<sup>-11</sup>] | 0% [0%, 0%] |  |  |

While univariate meta-regressions are informative, they can be confounded by covariation of the included variables. To account for this possibility, we also ran multivariate meta-regressions including the same variables used in the univariate analysis. Only comparisons with complete data for all variables were eligible for inclusion in the multivariate meta-regression. As a result, 571 comparisons were excluded, and 621 were retained for analysis. We then tested all 64 possible models, accounting for the combination of all variables. Table 6 shows the best model selected by the corrected Akaike Information Criteria (AICc), which includes Aβ aggregation status, Aβ concentration, and Aβ exposure duration and matches what was found by univariate meta-regressions. A similar multivariate meta-regression based on SMD was carried out, with results available in Supplementary Table 12. An additional analysis also indicated that the effects of Aβ concentration and exposure duration were synergistic, with toxicity becoming more dose-dependent as exposure duration increases (Supplementary Table 13).

**Table 6.** Three-level multivariate meta-regression model for testing selected protocol variables. Models have separate nested levels for articles and individual comparisons. The covariates tested were differentiation duration, Aβ state of aggregation, Aβ concentration, Aβ exposure duration, and cell density. Out of the 64 tested models, only the best fit, as selected by AICc, is shown (Akaike weight and AICc of 0.28 and 324.1, respectively). In the grey line are the sample size, intercept effect sizes (as absolute mean differences and cell viability) and p-value for the full range of moderators. White lines present betas indicating the additional contribution of each unit/category to the effect size and cell viability (as well as sample sizes and p-values for individual variables). I^2^, R^2^ and intercept p-value for the model were 99.4%, 19.3% and 0.087, respectively.

| Model and categories | Sample size | Effect size [95% C.I.] | Cell viability | p-value |
| --- | --- | --- | --- | --- |
| Aβ aggregation + Concentration + Exposure duration | 621 | -0.12 [-0.25, +0.02] | 88.7% [77.9%, 102%] | <0.0001 |
| Fibrils |  | -0.21 [-0.34, -0.07] | -18.9% [-28.8%, -6.8%] | 0.004 |
| Oligomers |  | -0.06 [-0.18, +0.07] | -5.8% [-16.5%, +7.3%] | 0.369 |
| Unclear |  | -0.14 [-0.27, -0.02] | -13.1% [-23.7%, -2%] | 0.026 |
| Aβ Concentration | | -1.2x10<sup>-2</sup> [-1.5x10<sup>-2</sup>, -1x10<sup>-2</sup>] | -1.2% [-1.5%, -1%] | <0.0001 |
| Exposure duration |  | -0.1 [-0.14, -0.05] | -9.5% [-13.1%, -4.9%] | <0.0001 |

## Discussion

The human neuroblastoma cell line SH-SY5Y has been extensively employed as a cell model to investigate the toxicity of Aβ, as indicated by the large number of published articles retrieved from our search. To the best of our knowledge, this is the first systematic review to synthesize published data on Aβ-induced toxicity in SH-SY5Y cells. Our findings include both a scoping review of methodological and reporting practices at the article level and a meta-analysis and meta-regression to identify factors that impact the observed toxicity outcomes. The evidence compiled in this review demonstrates that Aβ exposure reduces the viability of SH-SY5Y cells, as assessed by the MTT reduction assay, suggesting impairment of mitochondrial metabolic activity and, by extension, neuronal viability.

Metabolic-based viability assays were the most commonly used readouts, with MTT being the most widely employed reagent. In the MTT assay, tetrazolium salts are reduced by metabolically active cells to formazan, a water-insoluble violet-blue product that can be quantified by absorbance, which is assumed to be proportional to the number of metabolically viable cells (Mosmann, 1983). Although the MTT assay is widely used to infer cell proliferation, viability, or mitochondrial activity, its underlying mechanism is still not fully elucidated, and there are reports of MTT reduction through extramitochondrial enzymes (Berridge and Tan, 1993; Ghasemi et al., 2021). As a result, it has been proposed that the MTT assay should be interpreted primarily as a measure of the combined activity of enzymes involved in cellular metabolism, rather than as a direct indicator of cell viability (Berridge et al., 2005). In addition, variables such as medium composition and MTT reagent concentration may also contribute to the over- or underestimation of results (Surin et al., 2017; Ghasemi et al., 2021; Swain et al., 2025).

Approximately 70% of the studies included in our review employed SH-SY5Y cells to evaluate interventions aimed at counteracting Aβ-induced toxicity. This predominant use of a neuroprotection-focused framework may partly explain the lower-than-expected number of studies reporting preserved or increased cell viability following Aβ exposure, as observed in the funnel plot. Such outcomes may have been perceived as methodologically incongruent with the study objectives (as demonstrating neurotoxicity as a prerequisite for testing protective effects) and therefore deemed unpublishable or uninformative. This aligns with prior findings on publication bias in biomedical sciences (Fanelli, 2012).

Poor reporting quality significantly hindered the interpretation and synthesis of results across studies. Notably, the absence of standardized definitions for the experimental unit in *in vitro* research limited our ability to interpret reported sample sizes accurately. A wide range of ambiguous descriptions was used, often failing to clearly distinguish between technical and biological replicates. This lack of clarity may partly account for the unusually large standardized effect sizes observed in some comparisons, as technical replicates typically exhibit lower variability than biological replicates, potentially leading to inflated effect sizes when the former are incorrectly treated as independent observations. Additional factors that may contribute to these extreme effect sizes include inaccuracies in the extraction of means and error values from figures - particularly when error bars were small - as well as selective reporting practices (i.e., exclusion of outliers), and eventually data fabrication. These very large effect sizes were part of the reason why heterogeneity between results was very high across the sample.

Among the 359 articles from which data were extracted, 44.6% did not report the origin of the cell line used, and 4.7% stated that it was acquired by donation. Along with the absence of cell line authentication, this raises concerns about the reliability of the cell lines used. This is particularly relevant for cell lines like SH-SY5Y, which, being tumor-derived, are especially prone to phenotypic and genetic drift during prolonged culture (Fusenig et al., 2017). Furthermore, the risk of cross-contamination between cell lines remains a well-documented issue (Capes-Davis et al., 2010; Fusenig et al., 2017), and, without routine authentication procedures, such contamination may go undetected, especially when morphological differences are subtle or absent. These concerns, when considered alongside others, such as poor definition of experimental units, highlight the need for rigorous assessment of internal validity of *in vitro* studies (Whaley et al., 2024). The development and implementation of standardized tools to evaluate risk of bias in cell culture research is therefore critical for advancing evidence synthesis in this field, and ongoing initiatives are beginning to address this gap (Mathisen et al., 2025; Network et al., 2020; Tran et al., 2021).

In terms of reporting quality, a significant proportion of experiments failed to specify the source of Aβ or the species origin of its amino acid sequence. Although it is reasonable to assume the peptide used is usually synthetic and based on the human sequence, clearer reporting of these variables is recommended, as recombinantly produced peptides or sequences from rats were used in some studies. Likewise, almost half of the experiments did not provide a clear description of the aggregation state of Aβ and likely used heterogeneous mixtures of soluble and insoluble aggregates, which may impact toxicity identification. This description could be provided by employing conformational antibodies capable of discriminating different Aβ aggregates, such as A11 (Kayed et al., 2003; Glabe, 2004), NU1 (Lambert et al., 2007) and NUsc1 (Sebollela et al., 2017), selectives for oligomeric species, and OC, selective for pre-fibrillar species (Kayed et al., 2007).

This lack of proper description is concerning, given the differential toxicity of soluble and insoluble Aβ aggregates (Benilova et al., 2012; De et al., 2019; Figueiredo et al., 2013). The clinical implication of this differential role of distinct Aβ aggregates is underscored by findings reporting that amyloid plaque burden does not consistently correlate with cognitive decline in Alzheimer’s disease (Price & Morris, 1999; Selkoe & Hardy, 2016). It is therefore strongly recommended to carefully select and characterize Aβ aggregation status when using SH-SY5Y cells to evaluate novel therapeutic compounds for Aβ toxicity, with the ultimate goal of developing effective treatments for Alzheimer’s disease.

The mean Aβ concentration used across the studies included in this review was 41.5 μM, a high value compared to the estimated concentrations found *in vivo*. For instance, the levels of soluble Aβ oligomers found in brain tissue from AD patients have been reported to be in the low picogram per gram range (Mc Donald et al., 2010; Yang et al., 2013), and neurotoxicity has been induced *in vitro* by human brain-derived Aβ oligomers at concentrations as low as 50 nM (Gong et al., 2003). Interestingly, the median Aβ concentration in our review, 10 μM, was significantly lower than the mean, indicating a right-skewed distribution driven by a subset of studies using concentrations in the millimolar range. This discrepancy points to a potential lack of awareness regarding the low aqueous solubility of Aβ - particularly the 42-residue isoform - and highlights a broader issue of insufficient standardization in *in vitro* neurotoxicity models. Despite these caveats, our meta-analysis revealed a robust and statistically significant reduction in cell viability following Aβ exposure, consistent across both standard random-effects models and three-level models accounting for clustering at the study level.

Although a toxic effect was observed in the majority of studies, the size of this effect was very heterogeneous, both across studies and across experiments in the same study. High heterogeneity is a very common observation in pre-clinical systematic reviews such as ours (Voelkl et al., 2018; Rosso et al., 2022; Ineichen et al., 2024). The fact that it was particularly high within studies suggests that this is partly due to experiments in a wide range of conditions, some of which may function primarily as positive or negative controls. There was also substantial variability in the experimental conditions in which the model was employed across studies, including differences in cell culture conditions such as medium, supplements, number of passages, and Aβ concentration.

To evaluate whether experimental conditions could explain part of this heterogeneity, we performed meta-regressions using multiple protocol variables. Predictably, both Aβ concentration and exposure duration moderated the effect, with both variables having a synergistic effect in an interaction analysis, indicating that the dose-dependency of the effect increases with time. This suggests that prolonged exposure to low doses may produce effects that are comparable to those observed after shorter exposure to higher concentrations.

In contrast, differentiation status showed only a nonsignificant trend towards decreased toxicity with longer differentiation periods. This result was somewhat unexpected, given that Aβ toxicity has been shown to involve direct interactions with synaptic structures (Alfonso et al., 2014; Lacor et al., 2007; Marttinen et al., 2018), which are less abundant in undifferentiated SH-SY5Y cells than in differentiated ones (Kovalevich & Langford, 2013; Lopes et al., 2019). The observed trend may therefore be spurious or influenced by methodological heterogeneity. It is also possible that limiting our analysis to viability assays may have prevented us from observing more subtle signs of Aβ toxicity in differentiated cells. In line with this notion, individual studies have reported increased susceptibility to Aβ toxicity in differentiated SH-SY5Y cells relative to undifferentiated ones (Krishtal et al., 2019). The absence of a robust effect in our meta-analysis may reflect the influence of uncontrolled confounding variables or the possibility that, at supraphysiological Aβ concentrations, toxicity mechanisms may become less selective and less dependent on neuronal maturity.

We also examined the impact of Aβ aggregation state on cellular toxicity. Interestingly, compared to monomeric Aβ, which has been reported to be non-toxic and may even play physiological roles at low concentrations (Giuffrida et al., 2009; Jeong et al., 2022), both fibrillar Aβ and preparations of undefined aggregation status were associated with significantly higher toxicity. In contrast, Aβ preparations classified as oligomeric were associated with slightly lower toxicity, although this difference was not statistically significant. This finding was also unexpected, given the robust body of evidence pointing to soluble oligomers as the most neurotoxic Aβ species (Cline et al., 2018; Selkoe & Hardy, 2016; Tolar et al., 2021). As with differentiation status, we cannot rule out the possibility that poor reporting, inadequate biochemical characterization and classification of Aβ preparations, or other unidentified confounders may have biased this assessment.

Finally, a growing number of studies have reported that highly toxic Aβ assemblies exist along the continuum between soluble oligomers and insoluble fibrils. These species are distinct from typical protofibrils - an on-pathway to fibrils species - and are commonly found in close interaction with mature fibrils (Bigi et al., 2022; Lasagna-Reeves et al., 2011). It seems reasonable to consider that preparations categorized as fibrils, as well as those that were not characterized (“unclear”), may contain and release such toxic assemblies, in contrast to typical fibril-free oligomeric preparations. Therefore, the observed higher Aβ toxicity retrieved from studies using fibrils or unclear aggregation status could be instigated by intermediate species that are absent in well-characterized Aβ soluble oligomeric preparations. Future studies should prioritize rigorous biochemical characterization of Aβ aggregation states in order to clarify the contributions of specific assemblies to neurotoxicity in SH-SY5Y cell models.

## Limitations

A key limitation of this study stems from the absence of a standardized terminology to describe the experimental unit in the included studies. Despite our best efforts, it is possible that some technical replicates were misclassified as biological replicates. As outlined by Lazic et al. (2018), such misclassification can inflate the sample sizes and evidential strength of individual studies, potentially biasing the overall effect estimates (Lazic et al., 2018). While we sought to mitigate this risk by avoiding the use of standardized mean differences, studies with artificially precise data are still likely to weigh more in a meta-analysis. Additionally, our approach to variable categorization in the meta-regression, particularly regarding Aβ species, may have introduced misclassification bias. Beyond instances where the aggregation state was explicitly unreported, it is possible that other preparations were incorrectly categorized due to inconsistencies or limitations in how aggregation states were described. This lack of precise characterization may have affected the accuracy of subgroup comparisons and should be addressed in future primary studies through improved reporting and biochemical validation.

Additionally, restricting data extraction and analysis to metabolic-based viability assays also represents a limitation. Although these assays were the most commonly employed readout, they may fail to detect earlier and more sensitive toxicity signals compared with other methods, potentially limiting the scope of our conclusions. Furthermore, the small number of experiments with differentiated cells and variability between multiple protocol variables precluded the classification of differentiation protocols based on the differentiating agent, limiting our analysis to the presence of differentiation and its duration.

Another limitation of this study relates to the time frame of data collection. Due to feasibility constraints and the extended duration of data extraction and analysis, our literature search included only articles published up to December 2020. A follow-up search using the original search terms, limited to articles published from December 2020 onward, retrieved an additional 3,202 articles in PubMed and 1,926 in Embase (see Supplementary Figure 4). These were screened using the same eligibility criteria with the support of an AI-based tool, resulting in the inclusion of 541 articles. Based on the inclusion rate observed in our review, we estimate that approximately 142 of these would have met our inclusion criteria, representing a potential increase of 39.6% in the number of included articles.

While we acknowledge this as a significant limitation, the large number of studies already included in our analysis and the robustness of the overall effect size suggest that the inclusion of more recent data would be unlikely to substantially alter our main conclusions. A scoping review of 10% of included articles by full-text analysis showed general article features to be similar to those from the original sample, including the predominance of cell viability assays (76%), Aβ_42_ (82%), oligomers as aggregation state (32%) and non-differentiated cells (86%) (Supplementary Table 9). This suggests that the general profile of included articles remains unaltered, and that the conclusions from our time-limited sample are likely to hold in more recent studies.

## Recommendations for future studies

Improved reporting of methodological details and clearer definitions of experimental units are essential for enhancing the transparency and reproducibility of *in vitro* research. To address these issues, we recommend the adoption of standardized, unambiguous terminologies for the classification of replicates and experimental design. Lazic et al. (2018) advocate for replacing ambiguous terms such as ’technical’ and ’biological’ replicates with more precise descriptors - namely, biological, experimental, and observational units - which help clarify the hierarchical structure of the experimental setup (Lazic et al., 2018). In cell line studies, the use of standardized language specifying the variable components between experimental units (e.g., passage number, culture plate, or experimental day) would further aid in reproducibility and comparability across studies. Additionally, routine implementation of quality control measures - such as cell line authentication and mycoplasma testing - is strongly encouraged, as these practices enhance the reliability and translational relevance of findings derived from cell-based models. For systematic reviews in particular, the development and dissemination of tools and guidelines for assessing risk of bias in *in vitro* studies would represent a valuable step toward improving the rigor and interpretability of evidence synthesis in this field.

## Conclusions

Our meta-analysis confirms that Aβ exposure significantly reduces SH-SY5Y cell viability, as measured by the metabolic-based cell viability assays. The magnitude of this effect is primarily influenced by Aβ concentration, exposure duration, and aggregation state. In contrast, other factors such as differentiation status and cell density did not show consistent effects across studies. Our analysis also identified widespread methodological shortcomings, including inadequate reporting practices, lack of cell line authentication, and insufficient characterization of Aβ preparations. These limitations reduce the reliability and reproducibility of individual studies and may affect the interpretability of compiled findings. Overall, our findings support the continued use of SH-SY5Y cells as a relevant *in vitro* model for investigating Aβ-induced neurotoxicity. However, they also highlight the urgent need for improved methodological rigor, standardized reporting, and quality control measures to enhance the model’s reliability and translational value in AD research.

## Acknowledgements

The authors acknowledge the contributions of the BRISA Consortium members to the planning, conduct, and interpretation of the results of this study, and are particularly indebted to Francisco Marques da Silva Neto for assistance with LLM-based screening. They would also like to thank Wolfgang Viechtbauer for input on the analysis.

## Data Availability Statement

All data generated and analyzed during this study are included in the manuscript, supporting information or available at indicated databases.

## Author contributions

Conceptualization, methodology, project administration, resources, supervision, validation, and visualization of the study were carried out by NRP, CFDC, GSC, OBA, and AS. These authors were also responsible for the original draft preparation as well as the critical review and editing of the manuscript. Funding acquisition was secured by OBA and AS. Statistical analyses were performed mainly by CFDC, with contributions from NRP, GSC, and OBA. All authors contributed to data curation and investigation and reviewed the manuscript, collectively assuming responsibility for the decision to submit it for publication.

## List of abbreviations

AD: Alzheimer’s Disease
AICc: Corrected Akaike Information Criteria
APP: Amyloid Precursor Protein
ATRA: All Trans Retinoic Acid
ATCC: American Type Culture Collection
Aβ: Beta-amyloid peptide
BDNF: Brain-Derived Neurotrophic Factor
CI: Confidence Interval
DMEM: Dulbecco’s Modified Eagle Medium
ECACC: European Collection of Authenticated Cell Cultures
FBS: Fetal Bovine Serum
FCS: Fetal Calf Serum
FGF: Fibroblast growth factor
F12: Nutrient Mixture F12
MTT: 3-(4,5-dimethylthiazol-2-yl)-2,5-diphenyltetrazolium bromide
N: Number
N2: N-2 Supplement
ROM: Ratio of Means
SEM: Standard Error of the Mean
SD: Standard Deviation
SMD: Standardized Mean Difference

## References

1. Agholme, L., Lindström, T., Kågedal, K., Marcusson, J., & Hallbeck, M. (2010). An In Vitro Model for Neuroscience: Differentiation of SH-SY5Y Cells into Cells with Morphological and Biochemical Characteristics of Mature Neurons. Journal of Alzheimer’s Disease, 20(4), 1069–1082. 10.3233/JAD-2010-091363

2. Alfonso, S., Kessels, H. W., Banos, C. C., Chan, T. R., Lin, E. T., Kumaravel, G., Scannevin, R. H., Rhodes, K. J., Huganir, R., Guckian, K. M., Dunah, A. W., & Malinow, R. (2014). Synapto-depressive effects of amyloid beta require PICK 1. European Journal of Neuroscience, 39(7), 1225–1233. 10.1111/ejn.12499

3. Benilova, I., Karran, E., & De Strooper, B. (2012). The toxic Aβ oligomer and Alzheimer’s disease: An emperor in need of clothes. Nature Neuroscience, 15(3), 349–357. 10.1038/nn.3028

4. Berridge, M. V., Herst, P. M., & Tan, A. S. (2005). Tetrazolium dyes as tools in cell biology: New insights into their cellular reduction. In Biotechnology Annual Review (Vol. 11, pp. 127–152). Elsevier. 10.1016/S1387-2656(05)11004-7

5. Berridge, M. V., & Tan, A. S. (1993). Characterization of the Cellular Reduction of 3-(4,5-dimethylthiazol-2-yl)-2,5-diphenyltetrazolium bromide (MTT): Subcellular Localization, Substrate Dependence, and Involvement of Mitochondrial Electron Transport in MTT Reduction. Archives of Biochemistry and Biophysics, 303(2), 474–482. 10.1006/abbi.1993.1311

6. Biedler, J. L., Roffler-Tarlov, S., Schachner, M., & Freedman, L. S. (1978). Multiple Neurotransmitter Synthesis by Human Neuroblastoma Cell Lines and Clones. Cancer Research, 38.

7. Bigi, A., Cascella, R., Chiti, F., & Cecchi, C. (2022). Amyloid fibrils act as a reservoir of soluble oligomers, the main culprits in protein deposition diseases. BioEssays, 44(11), 2200086. 10.1002/bies.202200086

8. Burdick, D., Soreghan, B., Kwon, M., Kosmoski, J., Knauer, M., Henschen, A., Yates, J., Cotman, C., & Glabe, C. (1992). Assembly and aggregation properties of synthetic Alzheimer’s A4/beta amyloid peptide analogs. Journal of Biological Chemistry, 267(1), 546–554. 10.1016/s0021-9258(18)48529-8

9. Calcagno, V. (2020). glmulti: Model Selection and Multimodel Inference Made Easy. https://CRAN.R-project.org/package=glmulti

10. Capes-Davis, A., Theodosopoulos, G., Atkin, I., Drexler, H. G., Kohara, A., MacLeod, R. A. F., Masters, J. R., Nakamura, Y., Reid, Y. A., Reddel, R. R., & Freshney, R. I. (2010). Check your cultures! A list of cross-contaminated or misidentified cell lines. International Journal of Cancer, 127(1), 1–8. 10.1002/ijc.25242

11. Cline, E. N., Bicca, M. A., Viola, K. L., & Klein, W. L. (2018). The Amyloid-β Oligomer Hypothesis: Beginning of the Third Decade. Journal of Alzheimer’s Disease, 64(s1), S567–S610. 10.3233/JAD-179941

12. Cummings, J. L., Morstorf, T., & Zhong, K. (2014). Alzheimer’s disease drug-development pipeline: Few candidates, frequent failures. Alzheimer’s Research & Therapy, 6(4), 37. 10.1186/alzrt269

13. De Medeiros, L. M., De Bastiani, M. A., Rico, E. P., Schonhofen, P., Pfaffenseller, B., Wollenhaupt-Aguiar, B., Grun, L., Barbé-Tuana, F., Zimmer, E. R., Castro, M. A. A., Parsons, R. B., & Klamt, F. (2019). Cholinergic Differentiation of Human Neuroblastoma SH-SY5Y Cell Line and Its Potential Use as an In vitro Model for Alzheimer’s Disease Studies. Molecular Neurobiology, 56(11), 7355–7367. 10.1007/s12035-019-1605-3

14. De, S., Wirthensohn, D. C., Flagmeier, P., Hughes, C., Aprile, F. A., Ruggeri, F. S., Whiten, D. R., Emin, D., Xia, Z., Varela, J. A., Sormanni, P., Kundel, F., Knowles, T. P. J., Dobson, C. M., Bryant, C., Vendruscolo, M., & Klenerman, D. (2019). Different soluble aggregates of Aβ42 can give rise to cellular toxicity through different mechanisms. Nature Communications, 10(1), 1541. 10.1038/s41467-019-09477-3

15. Encinas, M., Iglesias, M., Liu, Y., Wang, H., Muhaisen, A., Ceña, V., Gallego, C., & Comella, J. X. (2000). Sequential Treatment of SH-SY5Y Cells with Retinoic Acid and Brain-Derived Neurotrophic Factor Gives Rise to Fully Differentiated, Neurotrophic Factor-Dependent, Human Neuron-Like Cells. Journal of Neurochemistry, 75(3), 991–1003. 10.1046/j.1471-4159.2000.0750991.x

16. Fanelli, D. (2012). Negative results are disappearing from most disciplines and countries. Scientometrics, 90(3), 891–904. 10.1007/s11192-011-0494-7

17. Figueiredo, C. P., Clarke, J. R., Ledo, J. H., Ribeiro, F. C., Costa, C. V., Melo, H. M., Mota-Sales, A. P., Saraiva, L. M., Klein, W. L., Sebollela, A., De Felice, F. G., & Ferreira, S. T. (2013). Memantine Rescues Transient Cognitive Impairment Caused by High-Molecular-Weight A Oligomers But Not the Persistent Impairment Induced by Low-Molecular-Weight Oligomers. Journal of Neuroscience, 33(23), 9626–9634. 10.1523/JNEUROSCI.0482-13.2013

18. Fontana, I. C., Zimmer, A. R., Rocha, A. S., Gosmann, G., Souza, D. O., Lourenco, M. V., Ferreira, S. T., & Zimmer, E. R. (2020). Amyloid-β oligomers in cellular models of Alzheimer’s disease. Journal of Neurochemistry, 155(4), 348–369. 10.1111/jnc.15030

19. Forster, J. I., Köglsberger, S., Trefois, C., Boyd, O., Baumuratov, A. S., Buck, L., Balling, R., & Antony, P. M. A. (2016). Characterization of Differentiated SH-SY5Y as Neuronal Screening Model Reveals Increased Oxidative Vulnerability. SLAS Discovery, 21(5), 496–509. 10.1177/1087057115625190

20. Fusenig, N. E., Capes-Davis, A., Bianchini, F., Sundell, S., & Lichter, P. (2017). The need for a worldwide consensus for cell line authentication: Experience implementing a mandatory requirement at the International Journal of Cancer. PLOS Biology, 15(4), e2001438. 10.1371/journal.pbio.2001438

21. Ghasemi, M., Turnbull, T., Sebastian, S., & Kempson, I. (2021). The MTT Assay: Utility, Limitations, Pitfalls, and Interpretation in Bulk and Single-Cell Analysis. International Journal of Molecular Sciences, 22(23), 12827. 10.3390/ijms222312827

22. Giuffrida, M. L., Caraci, F., Pignataro, B., Cataldo, S., De Bona, P., Bruno, V., Molinaro, G., Pappalardo, G., Messina, A., Palmigiano, A., Garozzo, D., Nicoletti, F., Rizzarelli, E., & Copani, A. (2009). β-Amyloid Monomers Are Neuroprotective. The Journal of Neuroscience, 29(34), 10582–10587. 10.1523/JNEUROSCI.1736-09.2009

23. Glabe, C. (2004). Conformation-dependent antibodies target diseases of protein misfolding. Trends in Biochemical Sciences, 29(10), 542–547. 10.1016/j.tibs.2004.08.009

24. Glenner, G. G., & Wong, C. W. (1984). Alzheimer’s disease: Initial report of the purification and characterization of a novel cerebrovascular amyloid protein. Biochemical and Biophysical Research Communications, 120(3), 885–890. 10.1016/S0006-291X(84)80190-4

25. Gong, Y., Chang, L., Viola, K. L., Lacor, P. N., Lambert, M. P., Finch, C. E., Krafft, G. A., & Klein, W. L. (2003). Alzheimer’s disease-affected brain: Presence of oligomeric Aβ ligands (ADDLs) suggests a molecular basis for reversible memory loss. Proceedings of the National Academy of Sciences, 100(18), 10417–10422. 10.1073/pnas.1834302100

26. Hardy, J., & Selkoe, D. J. (2002). The Amyloid Hypothesis of Alzheimer’s Disease: Progress and Problems on the Road to Therapeutics. Science, 297(5580), 353–356. 10.1126/science.1072994

27. Hu, X., Crick, S. L., Bu, G., Frieden, C., Pappu, R. V., & Lee, J.-M. (2009). Amyloid seeds formed by cellular uptake, concentration, and aggregation of the amyloid-beta peptide. Proceedings of the National Academy of Sciences, 106(48), 20324–20329. 10.1073/pnas.0911281106

28. Ineichen, B. V., Held, U., Salanti, G., Macleod, M. R., & Wever, K. E. (2024). Systematic review and meta-analysis of preclinical studies. Nature Reviews Methods Primers, 4(1), 72. 10.1038/s43586-024-00347-x

29. Jan, A., Hartley, D. M., & Lashuel, H. A. (2010). Preparation and characterization of toxic Aβ aggregates for structural and functional studies in Alzheimer’s disease research. Nature Protocols, 5(6), 1186–1209. 10.1038/nprot.2010.72

30. Jeong, H., Shin, H., Hong, S., & Kim, Y. (2022). Physiological Roles of Monomeric Amyloid-β and Implications for Alzheimer’s Disease Therapeutics. Experimental Neurobiology, 31(2), 65–88. 10.5607/en22004

31. Kayed, R., Head, E., Sarsoza, F., Saing, T., Cotman, C. W., Necula, M., Margol, L., Wu, J., Breydo, L., Thompson, J. L., Rasool, S., Gurlo, T., Butler, P., & Glabe, C. G. (2007). Fibril specific, conformation dependent antibodies recognize a generic epitope common to amyloid fibrils and fibrillar oligomers that is absent in prefibrillar oligomers. Molecular Neurodegeneration, 2, 18. 10.1186/1750-1326-2-18

32. Kayed, R., Head, E., Thompson, J. L., McIntire, T. M., Milton, S. C., Cotman, C. W., & Glabe, C. G. (2003). Common Structure of Soluble Amyloid Oligomers Implies Common Mechanism of Pathogenesis. Science, 300(5618), 486–489. 10.1126/science.1079469

33. Kossmeier, M., Tran, U. S., & Voracek, M. (2020). Power-Enhanced Funnel Plots for Meta-Analysis: The Sunset Funnel Plot. Zeitschrift Für Psychologie, 228(1), 43–49. 10.1027/2151-2604/a000392

34. Kovalevich, J., & Langford, D. (2013). Considerations for the Use of SH-SY5Y Neuroblastoma Cells in Neurobiology. In S. Amini & M. K. White (Eds.), Neuronal Cell Culture (Vol. 1078, pp. 9–21). Humana Press. 10.1007/978-1-62703-640-5_2

35. Krishtal, J., Bragina, O., Metsla, K., Palumaa, P., & Tõugu, V. (2017). In situ fibrillizing amyloid-beta 1-42 induces neurite degeneration and apoptosis of differentiated SH-SY5Y cells. PLOS ONE, 12(10), e0186636. 10.1371/journal.pone.0186636

36. Krishtal, J., Metsla, K., Bragina, O., Tõugu, V., & Palumaa, P. (2019). Toxicity of Amyloid-β Peptides Varies Depending on Differentiation Route of SH-SY5Y Cells. Journal of Alzheimer’s Disease, 71(3), 879–887. 10.3233/JAD-190705

37. Lacor, P. N., Buniel, M. C., Furlow, P. W., Sanz Clemente, A., Velasco, P. T., Wood, M., Viola, K. L., & Klein, W. L. (2007). Aβ Oligomer-Induced Aberrations in Synapse Composition, Shape, and Density Provide a Molecular Basis for Loss of Connectivity in Alzheimer’s Disease. The Journal of Neuroscience, 27(4), 796–807. 10.1523/JNEUROSCI.3501-06.2007

38. Lambert, M. P., Velasco, P. T., Chang, L., Viola, K. L., Fernandez, S., Lacor, P. N., Khuon, D., Gong, Y., Bigio, E. H., Shaw, P., De Felice, F. G., Krafft, G. A., & Klein, W. L. (2007). Monoclonal antibodies that target pathological assemblies of Aβ. Journal of Neurochemistry, 100(1), 23–35. 10.1111/j.1471-4159.2006.04157.x

39. Lasagna-Reeves, C. A., Glabe, C. G., & Kayed, R. (2011). Amyloid-β Annular Protofibrils Evade Fibrillar Fate in Alzheimer Disease Brain. Journal of Biological Chemistry, 286(25), 22122–22130. 10.1074/jbc.M111.236257

40. Lazic, S. E., Clarke-Williams, C. J., & Munafò, M. R. (2018). What exactly is ‘N’ in cell culture and animal experiments? PLOS Biology, 16(4), e2005282. 10.1371/journal.pbio.2005282

41. Lopes, G. S., Lico, D. T. P., Silva-Rocha, R., De Oliveira, R. R., Sebollela, A., Paçó-Larson, M. L., & Larson, R. E. (2019). A phylogenetically conserved hnRNP type A/B protein from squid brain. Neuroscience Letters, 696, 219–224. 10.1016/j.neulet.2019.01.002

42. Marttinen, M., Takalo, M., Natunen, T., Wittrahm, R., Gabbouj, S., Kemppainen, S., Leinonen, V., Tanila, H., Haapasalo, A., & Hiltunen, M. (2018). Molecular Mechanisms of Synaptotoxicity and Neuroinflammation in Alzheimer’s Disease. Frontiers in Neuroscience, 12, 963. 10.3389/fnins.2018.00963

43. Mathisen, G. H., Svendsen, C., Vist, G. E., Husøy, T., Ames, H. M., Bearth, A., Audebert, M., Bernhard, A., Beronius, A., Bruzell, E. M., Di Consiglio, E., Davenport, M., Druwe, I., Geci, R., Gundert-Remy, U., Hartung, T., Hoffmann, S., Hogberg, H. T., Hooijmans, C. R., … Whaley, P. (2025). Identification of concepts of importance for the assessment of internal validity of in vitro toxicology studies using a modified Delphi technique. Evidence-Based Toxicology, 3(1), 2551013. 10.1080/2833373X.2025.2551013

44. Mathisen, G. H., Vist, G. E., Whaley, P., White, R. A., Husøy, T., Ames, H. M., Beronius, A., Di Consiglio, E., Druwe, I., Hartung, T., Hoffmann, S., Hooijmans, C. R., Machera, K., Prieto, P., Robinson, J. F., Roggen, E., Rooney, A. A., Roth, N., Spilioti, E., … Svendsen, C. (2023). Protocol: Testing the performance of INVITES-IN, a tool for assessing the internal validity of in vitro studies. 10.5281/ZENODO.8315605

45. Mc Donald, J. M., Savva, G. M., Brayne, C., Welzel, A. T., Forster, G., Shankar, G. M., Selkoe, D. J., Ince, P. G., & Walsh, D. M. (2010). The presence of sodium dodecyl sulphate-stable Aβ dimers is strongly associated with Alzheimer-type dementia. Brain, 133(5), 1328–1341. 10.1093/brain/awq065

46. Mehta, D., Jackson, R., Paul, G., Shi, J., & Sabbagh, M. (2017). Why do trials for Alzheimer’s disease drugs keep failing? A discontinued drug perspective for 2010-2015. Expert Opinion on Investigational Drugs, 26(6), 735–739. 10.1080/13543784.2017.1323868

47. Michno, W., Blennow, K., Zetterberg, H., & Brinkmalm, G. (2021). Refining the amyloid β peptide and oligomer fingerprint ambiguities in Alzheimer’s disease: Mass spectrometric molecular characterization in brain, cerebrospinal fluid, blood, and plasma. Journal of Neurochemistry, 159(2), 234–257. 10.1111/jnc.15466

48. Mosmann, T. (1983). Rapid colorimetric assay for cellular growth and survival: Application to proliferation and cytotoxicity assays. Journal of Immunological Methods, 65(1–2), 55–63. 10.1016/0022-1759(83)90303-4

49. Ouzzani, M., Hammady, H., Fedorowicz, Z., & Elmagarmid, A. (2016). Rayyan—A web and mobile app for systematic reviews. Systematic Reviews, 5(1), 210. 10.1186/s13643-016-0384-4

50. Pedersen, A.-K., Pfeiffer, A., Karemore, G., Akimov, V., Bekker-Jensen, D. B., Blagoev, B., Francavilla, C., & Olsen, J. V. (2021). Proteomic investigation of Cbl and Cbl-b in neuroblastoma cell differentiation highlights roles for SHP-2 and CDK16. iScience, 24(4), 102321. 10.1016/j.isci.2021.102321

51. Price, J. L., & Morris, J. C. (1999). Tangles and plaques in nondemented aging and ?preclinical? Alzheimer’s disease. Annals of Neurology, 45(3), 358–368. 10.1002/1531-8249(199903)45:3%253C358::AID-ANA12%253E3.0.CO;2-X

52. Querfurth, H. W., & LaFerla, F. M. (2010). Alzheimer’s Disease. New England Journal of Medicine, 362(4), 329–344. 10.1056/NEJMra0909142

53. R Core Team. (2024). R: A Language and Environment for Statistical Computing. R Foundation for Statistical Computing. https://www.R-project.org/

54. Rosso, M., Wirz, R., Loretan, A. V., Sutter, N. A., Pereira Da Cunha, C. T., Jaric, I., Würbel, H., & Voelkl, B. (2022). Reliability of common mouse behavioural tests of anxiety: A systematic review and meta-analysis on the effects of anxiolytics. Neuroscience & Biobehavioral Reviews, 143, 104928. 10.1016/j.neubiorev.2022.104928

55. Sebollela, A., Cline, E. N., Popova, I., Luo, K., Sun, X., Ahn, J., Barcelos, M. A., Bezerra, V. N., Lyra E Silva, N. M., Patel, J., Pinheiro, N. R., Qin, L. A., Kamel, J. M., Weng, A., DiNunno, N., Bebenek, A. M., Velasco, P. T., Viola, K. L., Lacor, P. N., … Klein, W. L. (2017). A human scFv antibody that targets and neutralizes high molecular weight pathogenic amyloid-β oligomers. Journal of Neurochemistry, 142(6), 934–947. 10.1111/jnc.14118

56. Sebollela, A., Mustata, G.-M., Luo, K., Velasco, P. T., Viola, K. L., Cline, E. N., Shekhawat, G. S., Wilcox, K. C., Dravid, V. P., & Klein, W. L. (2014). Elucidating Molecular Mass and Shape of a Neurotoxic Aβ Oligomer. ACS Chemical Neuroscience, 5(12), 1238–1245. 10.1021/cn500156r

57. Sehar, U., Rawat, P., Reddy, A. P., Kopel, J., & Reddy, P. H. (2022). Amyloid Beta in Aging and Alzheimer’s Disease. International Journal of Molecular Sciences, 23(21), 12924. 10.3390/ijms232112924

58. Selkoe, D. J., & Hardy, J. (2016). The amyloid hypothesis of Alzheimer’s disease at 25 years. EMBO Molecular Medicine, 8(6), 595–608. 10.15252/emmm.201606210

59. Seubert, P., Vigo-Pelfrey, C., Esch, F., Lee, M., Dovey, H., Davis, D., Sinha, S., Schiossmacher, M., Whaley, J., Swindlehurst, C., McCormack, R., Wolfert, R., Selkoe, D., Lieberburg, I., & Schenk, D. (1992). Isolation and quantification of soluble Alzheimer’s β-peptide from biological fluids. Nature, 359(6393), 325–327. 10.1038/359325a0

60. Stockert, J. C., Horobin, R. W., Colombo, L. L., & Blázquez-Castro, A. (2018). Tetrazolium salts and formazan products in Cell Biology: Viability assessment, fluorescence imaging, and labeling perspectives. Acta Histochemica, 120(3), 159–167. 10.1016/j.acthis.2018.02.005

61. Surin, A. M., Sharipov, R. R., Krasil’nikova, I. A., Boyarkin, D. P., Lisina, O. Yu., Gorbacheva, L. R., Avetisyan, A. V., & Pinelis, V. G. (2017). Disruption of functional activity of mitochondria during MTT assay of viability of cultured neurons. Biochemistry (Moscow), 82(6), 737–749. 10.1134/S0006297917060104

62. Swain, S., Roberts, D. M., Chowdhry, S., Durbin, R., Boyd, R., Petereit, J., & Renden, R. (2025). Low-Glucose Culture Conditions Bias Neuronal Energetics Towards Oxidative Phosphorylation. Journal of Neurochemistry, 169(6), e70125. 10.1111/jnc.70125

63. the Dominantly Inherited Alzheimer Network, Barthélemy, N. R., Li, Y., Joseph-Mathurin, N., Gordon, B. A., Hassenstab, J., Benzinger, Tammie, L. S., Buckles, V., Fagan, A. M., Perrin, R. J., Goate, A. M., Morris, J. C., Karch, C. M., Xiong, C., Allegri, R., Mendez, P. C., Berman, S. B., Ikeuchi, T., Mori, H., … McDade, E. (2020). A soluble phosphorylated tau signature links tau, amyloid and the evolution of stages of dominantly inherited Alzheimer’s disease. Nature Medicine, 26(3), 398–407. 10.1038/s41591-020-0781-z

64. Tolar, M., Hey, J., Power, A., & Abushakra, S. (2021). Neurotoxic Soluble Amyloid Oligomers Drive Alzheimer’s Pathogenesis and Represent a Clinically Validated Target for Slowing Disease Progression. International Journal of Molecular Sciences, 22(12), 6355. 10.3390/ijms22126355

65. Tran, L., Tam, D. N. H., Elshafay, A., Dang, T., Hirayama, K., & Huy, N. T. (2021). Quality assessment tools used in systematic reviews of in vitro studies: A systematic review. BMC Medical Research Methodology, 21(1), 101. 10.1186/s12874-021-01295-w

66. Velasco, P. T., Heffern, M. C., Sebollela, A., Popova, I. A., Lacor, P. N., Lee, K. B., Sun, X., Tiano, B. N., Viola, K. L., Eckermann, A. L., Meade, T. J., & Klein, W. L. (2012). Synapse-Binding Subpopulations of Aβ Oligomers Sensitive to Peptide Assembly Blockers and scFv Antibodies. ACS Chemical Neuroscience, 3(11), 972–981. 10.1021/cn300122k

67. Viechtbauer, W. (2010). Conducting Meta-Analyses in R with the metafor Package. Journal of Statistical Software, 36(3). 10.18637/jss.v036.i03

68. Voelkl, B., Vogt, L., Sena, E. S., & Würbel, H. (2018). Reproducibility of preclinical animal research improves with heterogeneity of study samples. PLOS Biology, 16(2), e2003693. 10.1371/journal.pbio.2003693

69. Walsh, D. M., Lomakin, A., Benedek, G. B., Condron, M. M., & Teplow, D. B. (1997). Amyloid β-Protein Fibrillogenesis. Journal of Biological Chemistry, 272(35), 22364–22372. 10.1074/jbc.272.35.22364

70. Whaley, P., Blain, R. B., Draper, D., Rooney, A. A., Walker, V. R., Wattam, S., Wright, R., & Hooijmans, C. R. (2024). Identifying assessment criteria for in vitro studies: A method and item bank. Toxicological Sciences, 201(2), 240–253. 10.1093/toxsci/kfae083

71. Wickham, H., Averick, M., Bryan, J., Chang, W., McGowan, L., François, R., Grolemund, G., Hayes, A., Henry, L., Hester, J., Kuhn, M., Pedersen, T., Miller, E., Bache, S., Müller, K., Ooms, J., Robinson, D., Seidel, D., Spinu, V., … Yutani, H. (2019). Welcome to the Tidyverse. Journal of Open Source Software, 4(43), 1686. 10.21105/joss.01686

72. Wickham, H., & Bryan, J. (2023). R Packages: Organize, Test, Document, and Share Your Code. O’Reilly Media. https://books.google.com.br/books?id=kTHFEAAAQBAJ

73. Yang, T., Hong, S., O’Malley, T., Sperling, R. A., Walsh, D. M., & Selkoe, D. J. (2013). New ELISAs with high specificity for soluble oligomers of amyloid β-protein detect natural Aβ oligomers in human brain but not CSF. Alzheimer’s & Dementia, 9(2), 99–112. 10.1016/j.jalz.2012.11.005

