## Supplementary information for "The human neuroblastoma SH-SY5Y cell line as a model to assess β-amyloid neurotoxicity: A systematic review and meta-analysis"

[a]Department of Biochemistry and Immunology, Ribeirao Preto Medical School, University of Sao Paulo, Ribeirao Preto, SP, Brazil.
[b]Institute of Medical Biochemistry Leopoldo de Meis, Federal University of Rio de Janeiro, Rio de Janeiro, RJ, Brazil.
[c]Institute of Biological Sciences, Federal University of Amazonas, Manaus, AM, Brazil.
[d]Mountain Research Center, Campus of Santa Apolónia, Polytechnic Institute of Bragança, Bragança, Portugal.
[e]Laboratory for Sustainability and Technology in Mountain Regions, Campus of Santa Apolónia, Polytechnic Institute of Bragança, Portugal.
[f]Brazilian Biosciences National Laboratory (LNBio), Brazilian Centre for Research in Energy and Materials (CNPEM), SP, Brazil.
[g]Laboratório de Produtos Bioativos, Federal University of Rio de Janeiro, Campus Macaé, Macaé, RJ, Brazil.
[h]Department of Medicine, University of Pernambuco, Garanhuns, PE, Brazil.
[i]Brazilian Reproducibility Initiative in preclinical Systematic review and meta-Analysis (BRISA), Brazil.

** These authors contributed equally to this work*

*# Corresponding authors:*
Adriano Sebollela


Olavo B. Amaral


*† present affiliation:*
MUSC, USA

**Supplementary information**

*Supplementary tables*

**Supplementary Table 1. Sentinel articles used to guide search strategy development.**

| Title | DOI/PMID | Publication year |
|---|---|---|
| Toxicity of Amyloid-β Peptides Varies Depending on Differentiation Route of SH-SY5Y Cells | 10.3233/JAD-190705 | 2019 |
| Cholinergic Differentiation of Human Neuroblastoma SH-SY5Y Cell Line and Its Potential Use as an In vitro Model for Alzheimer Disease Studies | 10.1007/s12035-019-1605-3 | 2019 |
| In situ fibrillizing amyloid-beta 1-42 induces neurite degeneration and apoptosis of differentiated SH-SY5Y cells | 10.1371/journal.pone.0186636 | 2017 |
| Autophagy Activation Alleviates Amyloid-β-Induced Oxidative Stress, Apoptosis and Neurotoxicity in Human Neuroblastoma SH-SY5Y Cells | 10.1007/s12640-017-9746-5 | 2017 |
| A critical role for the self-assembly of Amyloid-β1-42 in neurodegeneration | 10.1038/srep30182 | 2016 |
| Binding affinity of amyloid oligomers to cellular membranes is a generic indicator of cellular dysfunction in protein misfolding diseases | 10.1038/srep32721 | 2016 |
| Mitogen-activated protein kinase signaling pathways are involved in regulating α7 nicotinic acetylcholine receptor-mediated amyloid-β uptake in SH-SY5Y cells | 10.1016/j.neuroscience.2014.08.013 | 2014 |
| Intracellular localization of amyloid-β peptide in SH-SY5Y neuroblastoma cells | 10.3233/JAD-122455 | 2013 |
| Guanosine protects human neuroblastoma cells from oxidative stress and toxicity induced by Amyloid-beta peptide oligomers | 20846477 | 2010 |
| Differentiation increases the resistance of neuronal cells to amyloid toxicity | 10.1007/s11064-008-9627-7 | 2008 |
| Anti-oligomeric Abeta single-chain variable domain antibody blocks Abeta-induced toxicity against human neuroblastoma cells | 10.1016/j.jmb.2008.09.068 | 2008 |
| Calcium dysregulation and membrane disruption as a ubiquitous neurotoxic mechanism of soluble amyloid oligomers | 10.1074/jbc.M500997200 | 2005 |
| Oligomerization and toxicity of beta-amyloid-42 implicated in Alzheimer's disease | 10.1006/bbrc.2000.3051 | 2000 |
| β-Amyloid induces apoptosis in human-derived neurotypic SH-SY5Y cells | 10.1016/S0006-8993(96)00733-0 | 1996 |

**Supplementary Table 2. Search strings and numbers of retrieved articles in each database.**

| Database | Search string | Articles retrieved | Search date | Sentinel articles retrieved |
|---|---|---|---|---|
| PubMed | ("SH-SY5Y"[Title/Abstract] OR "SHSY5Y"[Title/Abstract] OR "SHSY-5Y"[Title/Abstract] OR "NEUROBLASTOMA"[Title/Abstract] OR "CELL LINE"[Title/Abstract] OR "neuronal cell*"[Title/Abstract]) AND ("AMYLOID*"[Title/Abstract] OR "ALZHEIMER"[Title/Abstract] OR "Abeta"[Title/Abstract] OR "betaA"[Title/Abstract] OR "Ass"[Title/Abstract] OR "ssA"[Title/Abstract]OR "βA"[Title/Abstract] OR "Aβ"[Title/Abstract]) | 8,710 | 21/10/2020 | 14 of 14 |
| Embase | ('sh-sy5y':ab,kw,ti OR 'shsy5y':ab,kw,ti OR 'shsy-5y':ab,kw,ti OR 'neuroblastoma':ab,kw,ti OR 'cell line':ab,kw,ti OR 'neuronal cell*':ab,kw,ti) AND ('alzheimer':ab,kw,ti OR 'amyloid*':ab,kw,ti OR 'abeta':ab,kw,ti OR 'betaa':ab,kw,ti OR 'aß':ab,kw,ti OR 'ßa':ab,kw,ti OR 'βa':ab,kw,ti OR 'aβ':ab,kw,ti) AND [article]/lim AND [english]/lim AND [embase]/lim | 7,237 | 29/10/2020 | 12 of 14 |

**Supplementary Table 3. Extracted data from included articles.**

| | |
|---|---|
| **Article** | DOI; first author; title; journal title; year of publication; whether interventions protecting against Aβ toxicity were tested; presence of a sample size calculation; presence of a conflict of interest statement |
| **Figure/Table** | Dependent variable; control group; experimental group |
| **Results** | Mean and SD (or SEM) for the control and treated groups in the plot; number of experimental units for the control and treated groups; experimental unit definition |
| **Protocol** | Toxicity assay; Aβ origin, Aβ isoform, Aβ aggregation state; Aβ species identity; duration and concentration of Aβ treatment; control description; cell source; whether cells were authenticated; whether they were tested for mycoplasma; number of cells plated per well; passage number; plate size; culture medium and supplements; whether cells were differentiated; differentiation medium, supplements and protocol |

**Supplementary Table 4. R packages used for analysis and visualization of data.**

| Packages | Version | RRIDs |
|---|---|---|
| Readxl | 1.4.3 | SCR_018083 |
| tidyverse | 2.0.0 | SCR_019186 |
| metafor | 4.6-0 | SCR_003450 |
| metaviz | 0.3.1 | SCR_025424 |
| glmulti | 1.0.8 | SCR_025423 |

**Supplementary Table 5. Toxicity assays used in the initial included comparisons.**

| Assay type | Comparisons | |
|---|---|---|
| | N | % |
| MTT or analogue reduction | 1386 | 54.0 |
| LDH release | 200 | 7.8 |
| Trypan blue exclusion test | 74 | 2.9 |
| Other cell viability assays | 184 | 7.2 |
| Direct indicators of cell death | 86 | 3.3 |
| Direct indicators of apoptosis | 296 | 11.5 |
| Other markers of cell death | 69 | 2.7 |
| Reactive oxygen species | 141 | 5.5 |
| Damage to mitochondrial membrane | 54 | 2.1 |
| Products of oxidative stress | 52 | 2.0 |
| Other oxidative stress assays | 19 | 0.7 |
| Unclear/Not recorded | 4 | 0.1 |

The percentage is given in relation to the 2565 included comparisons.

**Supplementary Table 6. Substrates used in metabolic viability-based assays.**

| Assay | Articles | |
|---|---|---|
| | N | % |
| MTT | 299 | 83.3 |
| WST | 23 | 6.4 |
| CCK-8 | 12 | 3.3 |
| MTS | 17 | 4.7 |
| XTT | 5 | 1.4 |
| EZ4U | 2 | 0.6 |
| Resazurin | 1 | 0.3 |

For comparisons, the percentage is given in relation to the 1192 included comparisons; for articles, the percentage is given in relation to 364 included articles. Abbreviations: MTT - 3-(4,5-dimethylthiazol-2-yl)-2,5-diphenyltetrazolium bromide; WST - Water Soluble Tetrazolium salts; CCK8 - Cell Counting Kit - 8; MTS - 3-(4,5-dimethylthiazol-2-yl)-5-(3-carboxymethoxyphenyl)-2-(4-sulfophenyl)-2H-tetrazolium; XTT - 2,3-bis-(2-methoxy-4-nitro-5-sulfophenyl)-2H-tetrazolium-5-carboxanilide; EZ4U - Easy For You.

**Supplementary Table 7. List of cell banks reported in the selected articles.**

| Cell bank | **N** | **%** |
|---|---|---|
| American Type Culture Collection (ATCC) | 109 | 30.4 |
| European Collection of Authenticated Cell Cultures (ECACC) | 30 | 8.4 |
| Chinese Academy of Sciences | 15 | 4.2 |
| Leibniz Institute DSMZ - German Collection of Microorganisms and Cell Cultures GmbH | 7 | 1.9 |
| National Centre for Cell Science (NCCS) | 5 | 1.4 |
| Korean Cell Line Bank | 3 | 0.8 |
| Riken Cell Bank | 3 | 0.8 |
| Pasteur Institute of Iran | 2 | 0.6 |
| Sigma-Aldrich | 2 | 0.6 |
| Institute of Biochemistry and Cell Biology | 1 | 0.3 |
| Invitrogen | 1 | 0.3 |
| LGC Promo-chem | 1 | 0.3 |
| The Cell Resource Centre of Institute of Basic Medicine | 1 | 0.3 |
| Zhong Qiao Xin Zhou Biotec Co., Ltd (Shanghai, China) | 1 | 0.3 |
| Not informed | 160 | 44.6 |

N refers to the number of articles, while the percentage is in relation to the 359 included articles.

**Supplementary Table 8. Cell culture medium and supplements used to maintain SH-SY5Y cells.**

| Protocol variable | Category | | Comparisons | |
|---|---|---|---|---|
| | | | N | % |
| Medium | DMEM_F12 | | 439 | 36.8 |
| | DMEM | | 375 | 31.5 |
| | MEM_F12 | | 118 | 9.9 |
| | MEM | | 56 | 4.7 |
| | RPMI | | 47 | 3.9 |
| | F12 | | 24 | 2.0 |
| | EMEM_F12 | | 21 | 1.8 |
| | EMEM | | 8 | 0.7 |
| | OptiMEM | | 3 | 0.3 |
| | Unclear | | 101 | 8.5 |
| Antibiotics | Yes | | 858 | 72 |
| | No | | 16 | 1.3 |
| | Unclear | | 318 | 26.7 |
| Glutamine | Yes | | 472 | 39.6 |
| | No | | 460 | 38.6 |
| | Unclear | | 260 | 21.8 |
| Serum type and concentration range | FBS/FCS | 6-10% | 761 | 63.8 |
| | | 11-15% | 206 | 17.3 |
| | | 16-20% | 17 | 1.4 |
| | | 1-5% | 11 | 0.9 |
| | | Not reported | 8 | 0.7 |
| | CS | 10% | 2 | 0.2 |
| | Combination | 5-10% | 5 | 0.4 |
| | No serum | - | 1 | 0.1 |
| | Not reported | Not reported | 181 | 15.2 |

Percentage corresponds to a fraction in relation to the 1192 included comparisons. Abbreviations: DMEM - Dulbecco's Modified Eagle Medium; F12 - Nutrient Mixture F12; MEM - Minimum Essential Medium; RPMI - Roswell Park Memorial Institute; EMEM - Eagle's Minimum Essential Medium; OptiMEM - Optimized Minimal Essential Medium; FBS - Fetal Bovine Serum; FCS - Fetal Calf Serum (synonym of FBS); CS - Calf Serum.

**Supplementary Table 9. Assay type and protocol variables from a contemporary sample (December 2020 to April 2025).**

| **Assay type** | | **N** | **%** |
|---|---|---|---|
| MTT or analogue reduction | | 22 | 75.9 |
| Other cell viability assays | | 8 | 27.6 |
| Markers of cell death and apoptosis | | 9 | 31 |
| Markers of oxidative stress | | 13 | 44.8 |
| **Protocol variable** | **Category** | **N** | **%** |
| Aβ isoform | 1-42 | 18 | 81.8 |
| | 1-40 | 2 | 9.1 |
| | Not reported | 2 | 9.1 |
| Aβ origin | Synthetic | 9 | 40.9 |
| | Recombinant | 1 | 4.5 |
| | Not reported | 12 | 54.5 |
| Aβ species identity | Human | 5 | 22.7 |
| | Not reported | 17 | 77.3 |
| Aβ aggregation | Oligomers | 7 | 18.2 |
| | Fibrils | 3 | 31.8 |
| | Monomers | 4 | 13.6 |
| | Not reported | 13 | 59.1 |
| Differentiation | Yes | 3 | 13.6 |
| | No | 19 | 86.4 |

The percentages for each assay type and protocol variable are based on the 29 articles included in the full-text screening and on 22 articles that use MTT or analog reduction assays, respectively.

**Supplementary Table 10. Trim-and-fill analysis.**

| **Method** | **Estimated number of missing studies on the right side** | **p-value** | **Adjusted meta-analysis** |
|---|---|---|---|
| R0 | 86 | <0.0001 | -0.40 [-0.42, -0.37], p<0.0001 |
| L0 | 0 | - | -0.46 [-0.48, -0.44], p<0.0001 |

The p-value for the null-hypothesis of no studies missing on the right side can only be obtained for method R0. The adjusted meta-analysis results include the effect size [95%C.I.] and the meta-analysis p-value.

**Supplementary Table 11. Three-level meta-regression models for protocol variables.**

| Moderators and categories | Sample size | Effect size [95% C.I.] | $I^2$ | $R^2$ | p-value |
|---|---|---|---|---|---|
| Duration of differentiation | 1144 | -6.2 [-6.6, -5.8] | 93.5% | 9.3% | 0.0134 |
| (day) | | 0.18 [0.04, 0.32] | | | 0.0134 |
| State of aggregation (reference=monomers) | 1192 (92) | -4.3 [-5.5, -3] | 94% | 1.4% | 0.0001 |
| Fibrils | 103 | -2.7 [-4.1, -1.2] | | | 0.0004 |
| Oligomers | 462 | -1.2 [-2.5, 0.1] | | | 0.0676 |
| Unclear | 535 | -2.4 [-3.8, -1] | | | 0.0006 |
| Concentration of Aβ | 1123 | -4.7 [-5.2, -4.2] | 99.6% | 10.2% | <0.0001 |
| (μM) | | -0.1 [-0.2, -0.1] | | | <0.0001 |
| Aβ exposure duration | 1180 | -4.9 [-5.7, -4.1] | 94.1% | 0% | 0.0002 |
| (day) | | -0.9 [-1.4, -0.4] | | | 0.0002 |
| Cell density | 690 | -6.4 [-7, -5.8] | 94.2% | 0% | 0.7549 |
| $10^3$Cells/cm$^2$ | | $7.5x10^{-11}$ [$4x10^{-10}$, $5.5x10^{-10}$] | | | 0.7549 |

Models have separate nested levels for articles and individual comparisons. Grey lines contain sample sizes, intercept effect sizes as ratio of means and cell viability, $R^2$ for the moderator and Q-test p-values for the moderator. White lines contain betas indicating the additional contribution of each unit/category to the effect size (as well as sample sizes and p-values for individual categories). For categorical variables, reference groups are described in the first column, with the sample size for these groups indicated in parentheses in the second column. P-values for all intercepts were <0.0001. Moderator $R^2$ values are calculated as the difference between total variances in the model with no moderators and in the tested model, divided by the total variance in the model with no moderators. Aβ concentrations above 100 μm were excluded from the concentration meta-regression.

**Supplementary Table 12. Three-level multivariate meta-regression model for testing selected protocol variables.**

| Model and categories | Sample size | Effect size [95% C.I.] | p-value |
|---|---|---|---|
| Aβ aggregation + Concentration + Exposure duration | 621 | -1.1 [-2.7, +0.5] | <0.0001 |
| Fibrils | | -2.1 [-3.6, -0.6] | 0.006 |
| Oligomers | | -1.2 [-2.6, +0.2] | 0.085 |
| Unclear | | -2.9 [-4.5, -1.4] | 0.0002 |
| Aβ Concentration | | -0.11 [-0.14, -0.09] | <0.0001 |
| Exposure duration | | -1.2 [-1.8, -0.6] | <0.0001 |

Models have separate nested levels for articles and individual comparisons. The covariates tested were differentiation duration, Aβ state of aggregation, Aβ concentration, Aβ exposure duration, and cell density. Out of the 64 tested models, only the best fit as selected by AICc is shown (Akaike weight and AICc of 0.24 and 3724.5, respectively). In the grey line are the sample size, intercept effect sizes as absolute mean differences and cell viability, p-value, and $R^2$ value for the full range of moderators. White lines present betas indicating the additional contribution of each unit/category to the effect size and cell viability (as well as sample sizes and p-values for individual variables). $I^2$, $R^2$ and intercept p-value for the model were 99.4%, 10.8% and 0.177, respectively.

**Supplementary Table 13. Interaction of Aβ concentration and exposure duration on cell viability.**

| Moderators and categories | Sample size | Effect size [95% C.I.] | Cell Viability | $R^2$ | p-value |
|---|---|---|---|---|---|
| No interaction | 1113 | -0.2 [-0.3, -0.17] | 80% [75.6%, 84.7%] | 16.6% | <0.0001 |
| Aβ concentration | | -0.013 [-0.015, -0.011] | -18.7% [-22.9%, -14.2%] | | <0.0001 |
| Aβ exposure duration | | -0.1 [-0.13, -0.06] | -10.9% [-12.4%, -9.3%] | | <0.0001 |
| Interaction | 1113 | -0.27 [-0.34, -0.2] | 76% [71%, 81.5%] | 16.6% | <0.0001 |
| Aβ concentration | | -0.008 [-0.01, -0.004] | -23.2% [-27.9%, -18.15%] | | 0.0002 |
| Aβ exposure duration | | -0.06 [-0.1, -0.01] | -18.5% [-19.5%, -17.3%] | | 0.01 |
| Aβ concentration + exposure duration | | -0.004 [-0.006, -0.001] | -23.6% [-28.5%, -18.4%] | | 0.008 |

Meta-regression analyses evaluating the effect of Aβ concentration and exposure time on cell viability, with or without considering the interaction between factors. Increasing Aβ concentration and longer exposure periods were associated with greater toxicity in both models. Models including the interaction indicated a negative interaction effect, suggesting that the impact of Aβ concentration increases with longer exposure times. Effect sizes are presented with 95% confidence intervals, and predicted cell viability values represent the expected biological response under each condition. $I^2$ and intercept p-values for all moderators were ~99.6% and <0.0001, respectively.

*Supplementary figures*

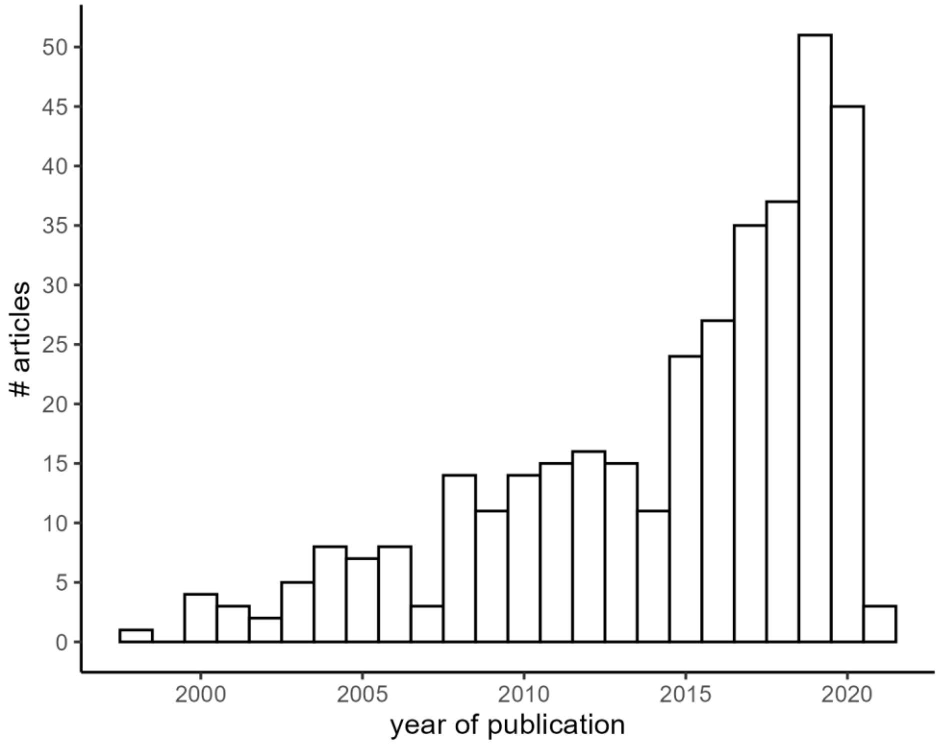


**Supplementary Figure 1. Year of publication of extracted articles.** Number of selected articles per year of publication, ranging from 1998 to 2021.

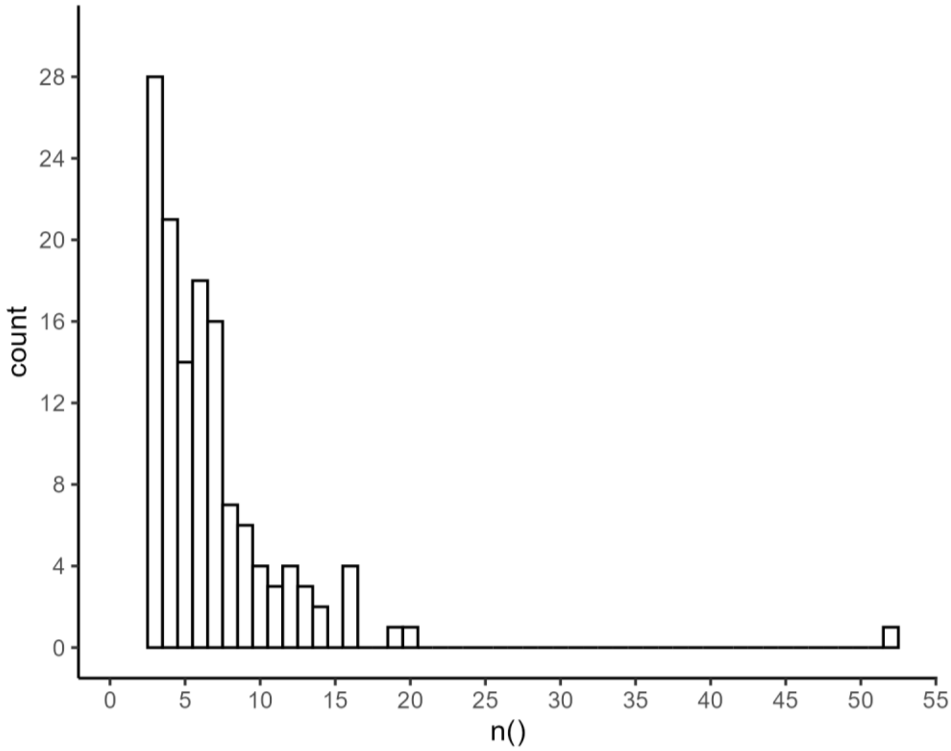


**Supplementary Figure 2. Number of comparisons per article.** The x-axis indicates the number of comparisons, and the y-axis the number of articles.

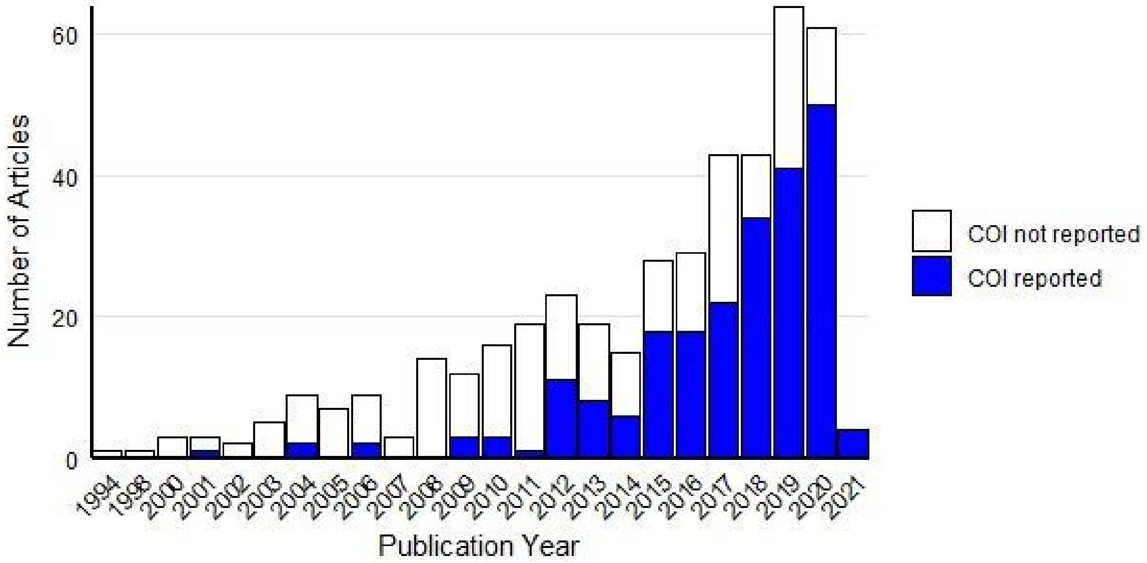


**Supplementary Figure 3. Number of articles per publication year that report a conflict of interest statement.** The x-axis indicates the publication year, and the y-axis the number of articles.

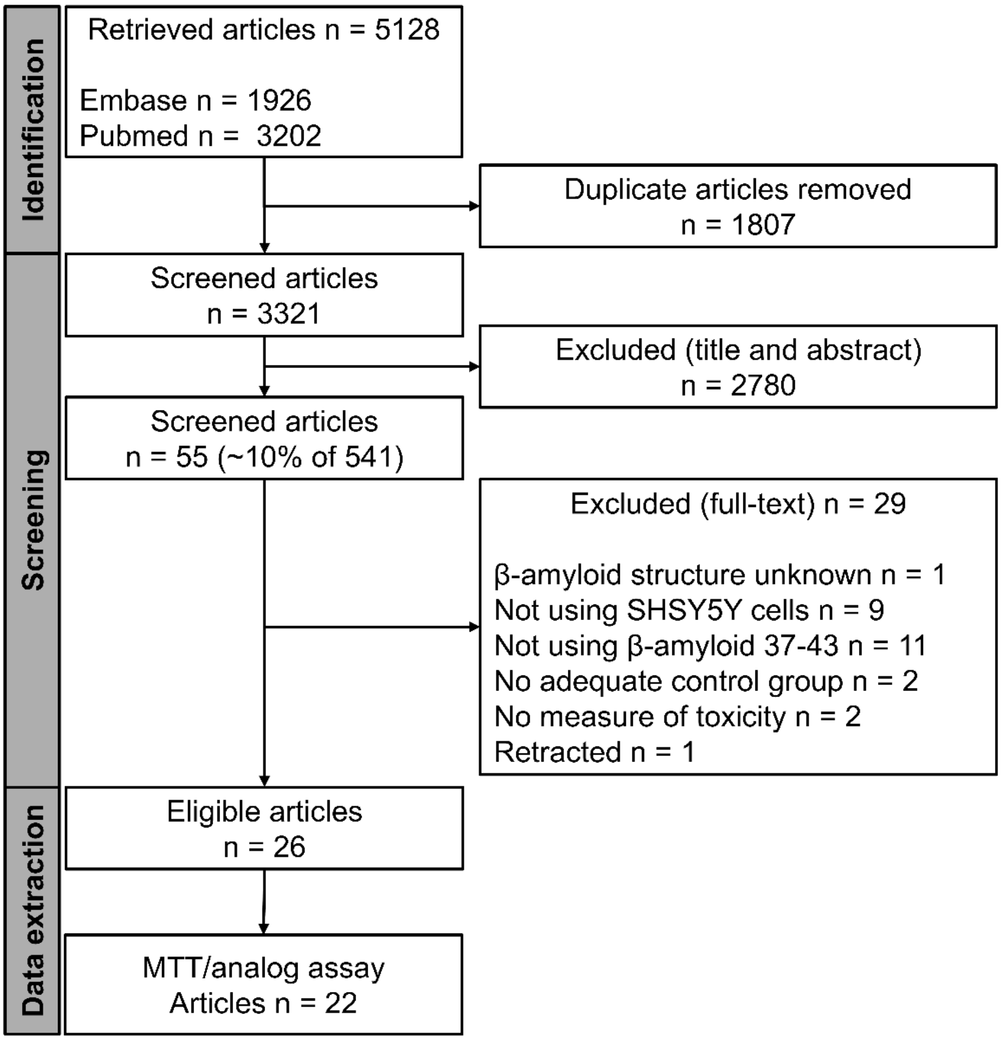


**Supplementary Figure 4. Flowchart of the scoping update.** Articles retrieved between December 2020 and April 2025, applying the same search strategy. After duplicate removal, 3,321 articles were retrieved. This set was subjected to an abstract screening using the same eligibility criteria with the support of an IA-based tool developed by our collaborators at BRISA (https://osf.io/fqe85/overview), resulting in the exclusion of 2,780 articles and the inclusion of 541. A full-text and qualitative data extraction was performed on a random sample of 10% of these articles. This step resulted in the exclusion of 26 articles and the inclusion of 29. Qualitative data were extracted from the 22 articles that employed MTT assay and analogs to assess Aβ-induced toxicity.

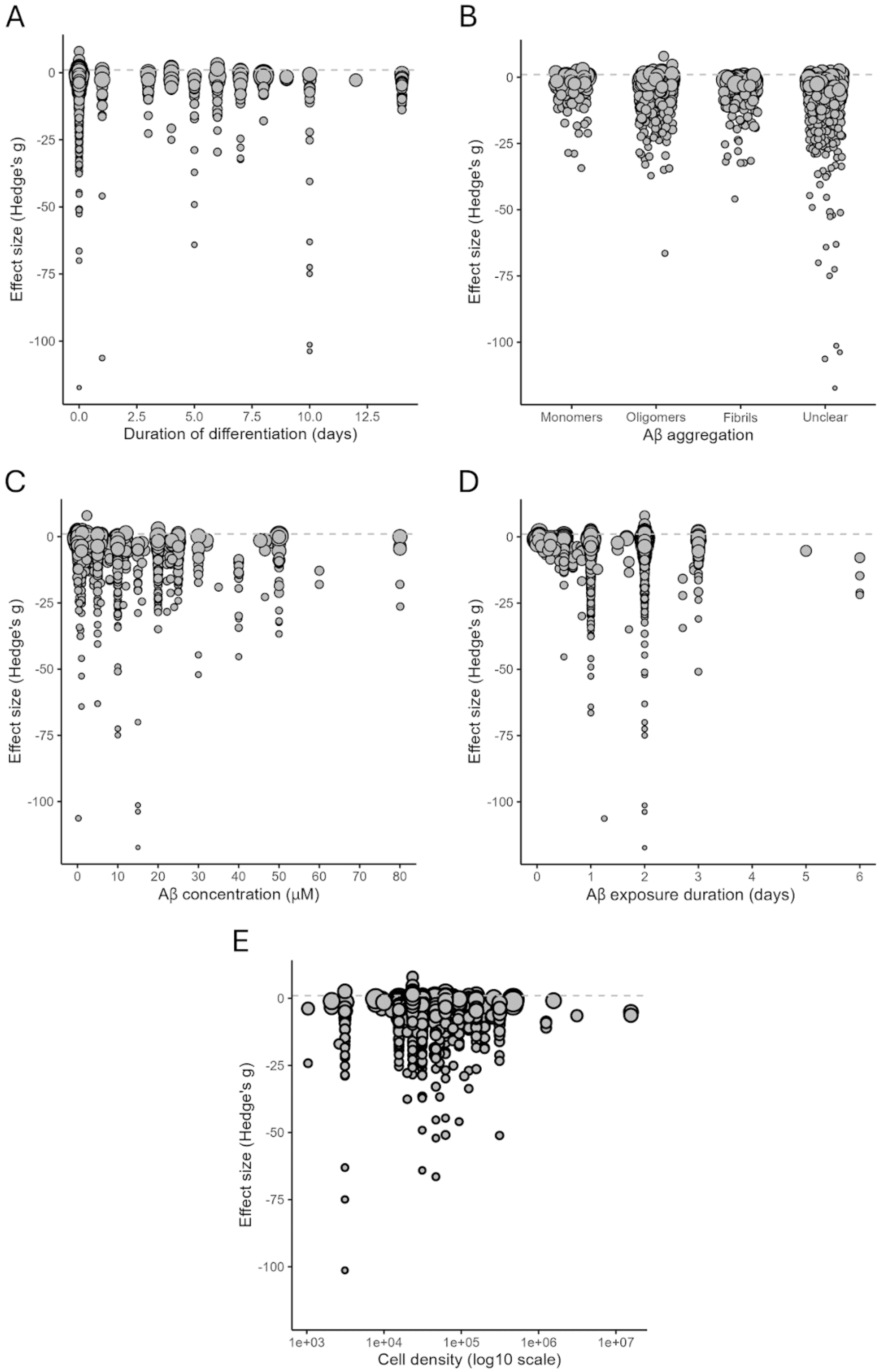


**Supplementary Figure 5. Univariate meta-regressions based on standardized mean differences as the effect size measure.** Each experiment is represented by an open circle, with size corresponding to the inverse variance (i.e., larger circles are more precise). Aβ concentrations above 100 μm were excluded from the concentration meta-regression.
